# The CovR regulatory network drives the evolution of Group B *Streptococcus* virulence

**DOI:** 10.1101/2021.01.25.428120

**Authors:** Maria-Vittoria Mazzuoli, Maëlle Daunesse, Hugo Varet, Isabelle Rosinski-Chupin, Rachel Legendre, Odile Sismeiro, Myriam Gominet, Pierre-Alexandre Kaminski, Philippe Glaser, Claudia Chica, Patrick Trieu-Cuot, Arnaud Firon

## Abstract

Virulence of the neonatal pathogen Group B *Streptococcus* depends on the master regulator CovR. Inactivation of CovR leads to large-scale transcriptome remodeling and impairs almost every step of the interaction between the pathogen and the host. However, comparative analyses suggested a plasticity of the CovR signalling pathway in clinical isolates, probably due to the host selective pressure and leading to phenotypic heterogeneity in the bacterial population. Here, we characterize the CovR regulatory network in a strain representative of the hypervirulent lineage responsible of the majority of late-onset meningitidis. Genome-wide binding and transcriptome analysis demonstrated that CovR acts as a direct and global repressor of virulence genes, either as a primary regulator or with specialized co-regulators. Remarkably, CovR directly regulates genes of the pan-genome, including the two specific hypervirulent adhesins and horizontally acquired genes, as well as core-genes showing mutational biases in the population. Parallel analysis of the CovR network in a second isolate links strain-specificities to micro-evolutions in CovR-regulated promoters and to broad difference due to variability in CovR activation by phosphorylation. Our results highlight the direct, coordinated, and strain-specific regulation of virulence genes by CovR. This intra-species evolution of the signalling network reshapes bacterial-host interactions, increasing the potential for adaptation and the emergence of clone associated with specific diseases.

## INTRODUCTION

*Streptococcus agalactiae*, commonly known as Group B *Streptococcus* (GBS), is the leading cause of sepsis and meningitis in the first three months of life and a significant cause of *in utero* infections and preterm births (1, 2). The reconstitution of the evolutionary history of the species suggests that the human-adapted strains have emerged in the mid-twentieth century, corresponding to the period of the first clinical cases (3). Of the five main clonal complexes (CCs) associated with human infections, strains of the CC-17 lineage are responsible of the vast majority of late-onset meningitis in neonates and, consequently, are classified as the hypervirulent GBS clones. CC-17 strains are specifically associated with human and are highly homogenous compared to strains belonging to other clonal complexes, a characteristic of an epidemic clone with worldwide dissemination (3–5).

The success of the hypervirulent clone as a neonatal pathogen is linked to the expression of two specific adhesins, HvgA and Srr2. The HvgA adhesin is a key determinant of the meningeal tropism of CC-17 strains by increasing GBS translocation across the blood-brain barrier (6), while the Srr2 serine-rich protein enhances the capacity of CC-17 strains to cross the intestinal barrier in the developing neonatal gastrointestinal tract (7, 8). Proteins covalently anchored to the cell-wall by their LPxTG motif through the activity of the sortase A enzyme (9), such as HvgA and Srr2, are a major group of virulence factors with adhesion or immune-modulatory properties (10). These cell-wall anchored proteins are subject to selective pressure generating variability in the GBS population either through allelic variation (*e*.*g* the *bibA/ hvgA* alleles) or gain and loss of virulence genes (*e*.*g* the *srr1/srr2* mutually exclusive loci) or mobile elements (10–12). In addition, a precise and coordinated control of the expression of virulence genes is essential for GBS pathogenesis. The expression of the appropriate combination of surface proteins and of secreted factors, most notably a ß-hemolytic/cytotoxic toxin (ß-h/c) (13, 14), is essential for GBS to establish commensal relationships within the adult vaginal and intestinal tracts as well as to become an extracellular pathogen in susceptible hosts (15–17).

The major regulator of host-pathogen interaction in ß-hemolytic streptococci is the transcriptional factor CovR belonging to the PhoP family of bacterial response regulator (18, 19). Targeted analysis in GBS demonstrated that CovR directly represses the transcription of the *cyl* operon encoding the ß-h/c toxin, the *bibA* gene, and the PI-1 pili operon (20–23). The regulation by CovR is highly dynamic and sustains the trade-off between cytotoxicity and adherence, ultimately leading to bacterial multiplication or elimination by the host immune response (13, 24, 25). The activity of CovR is modulated by its cognate histidine kinase CovS, which has a dual kinase and phosphatase activity on a conserved D_53_ aspartate residue (26, 27). The dynamic equilibrium between the opposite enzymatic activities depends on the interaction of CovS with the membrane protein Abx1 (26), and on the mutually exclusive phosphorylation of a CovR threonine residue (T_65_) by the serine-threonine kinase Stk1 (23).

In spite of being the major regulator of virulence in GBS, previous transcriptomic analyses of the CovR/S two-component system have been done in non-CC-17 isolates only. The inactivation of CovR is usually associated with a global transcriptome remodelling involving 10 to 15 % of the genes, with CovR being mainly a transcriptional repressor. However, important variation in the CovR regulon was observed early on among strains of different clonal complexes (21, 25, 26). In addition, a genomic analysis has highlighted mutational biases in the CovR/S system itself and in the CovR regulated promoters in CC-17 strains, suggestive of a positive selection acting on the CovR regulatory pathway in hypervirulent clones (4). Evolution of regulatory pathways is a common process observed between related bacterial species (28), but our knowledge on intra-species regulatory evolution remains limited (29–31). To address the role of CovR in GBS hypervirulent clones, we characterized the CovR regulon in a CC-17 strain and demonstrated the direct regulation of a combination of proteins involved in host-pathogen interaction. Comparative analysis revealed strain-specificities supported by mechanisms acting locally on CovR regulated genes and promoters and globally at the level of CovR activation by phosphorylation. The plasticity of the CovR regulatory network generates phenotypic heterogeneity at the species level, thus allowing the selection of clones associated to specific hosts and pathological conditions.

## RESULTS

### CovR regulates virulence genes expression and prophages transcription in BM110

To define the transcriptional response associated to CovR inactivation in the hypervirulent CC-17 lineage, we constructed two *covR* mutants in the BM110 strain (3). The first mutant has an in-frame deletion of the *covR* sequence (Δ*covR*) and the second a two base-pairs chromosomal substitution (AT->CC) resulting in the translation of a CovR_D53A_ variant that cannot be phosphorylated by CovS. Phenotypically, both mutants are hyper-haemolytic and hyper-pigmented, as observed for *covR* mutants in other backgrounds (Supplementary Fig. S1). Transcriptome analysis by RNA-seq of the CovR_D53A_ mutant revealed a differential expression for 12.2% of the genes (N = 266/2178; |Log_2_ FC| > 1; adjusted p-value < 0.005) at mid-exponential growth phase in rich medium (Supplementary Table S1A). Overall, fold changes associated to the 137 up-regulated genes were higher than those associated to the 129 down-regulated genes (Fig. 1A and Supplementary Tables S1B and S1C). Comparison of the CovR_D53A_ and Δ*covR* transcriptomes highlighted a clear correlation for highly differentially expressed genes but also emphasised specificities (Fig. 1B and 1C). The most striking difference was the opposite regulation of clusters of genes located in four prophages (Fig. 1D and Supplementary Fig. S2A). These four prophages accounted for a large proportion of the variability between mutants, encompassing 61.2% (79/129) of the down-regulated genes in the CovR_D53A_ and 34.2 % (128/374) of the up-regulated genes in the Δ*covR* mutant (Supplementary Tables S1).

**Figure 1.**
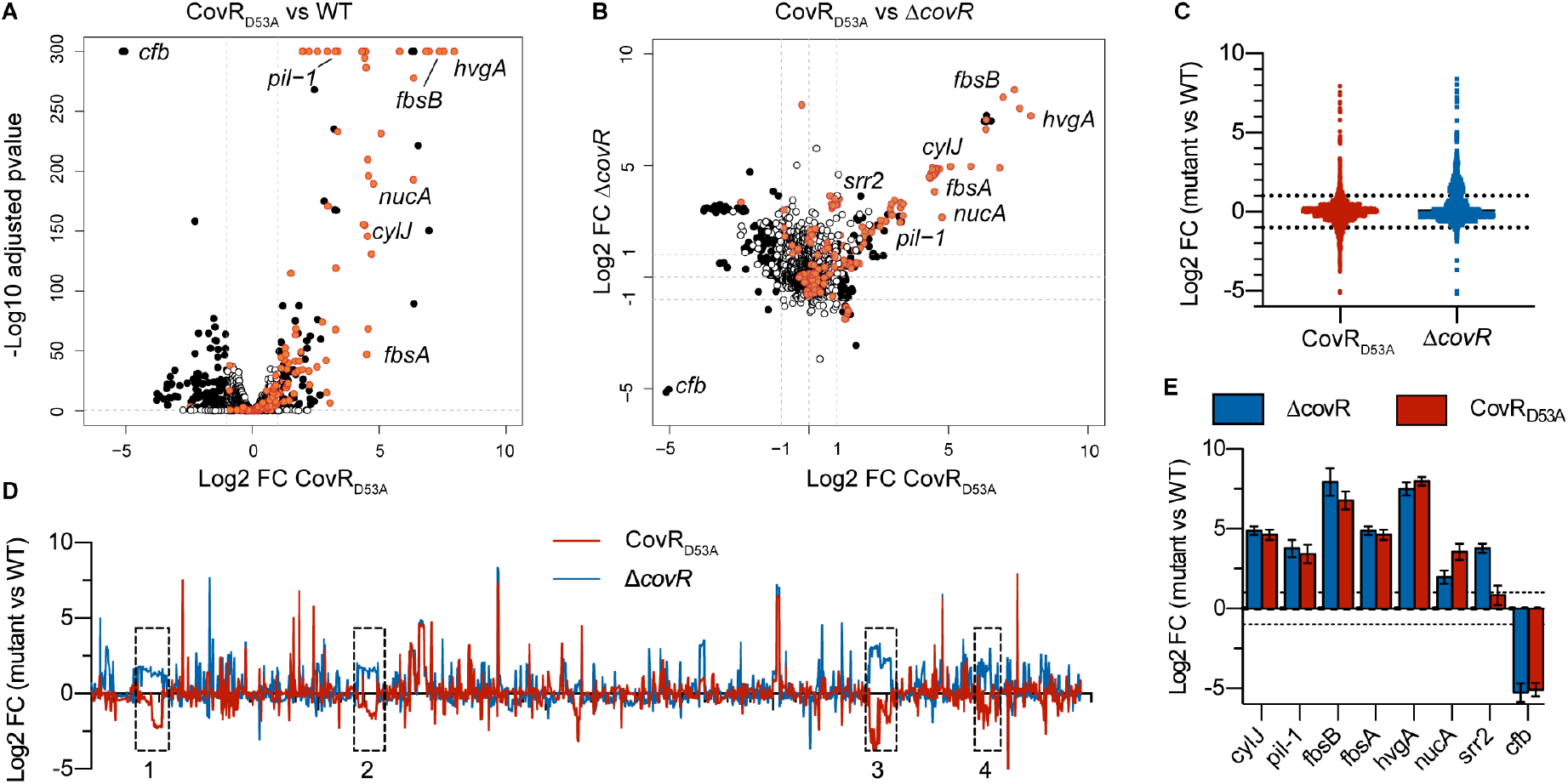
The CovR regulon in the hypervirulent BM110 strain. (A) Volcano plot of the BM110 CovR_D53A_ transcriptome at mid-exponential phase in THY. Each dot represents one of the 2178 genes with its RNA-seq fold change and adjusted p-value (Wald test) calculated from three independent replicates. Black and white dots symbolized significant (|Log_2_ FC| > 1; adjusted p-value < 0.005) and non-significant differentially transcribed genes, respectively. Red dots are genes associated to a CovR binding region identified by ChIP-Seq (see Fig. 2), with selected gene names highlighted. (B) Pairwise comparisons of the BM110 CovR_D53A_ and Δ*covR* mutant transcriptomes. Each dot has the same color-coded as in (A), corresponding to significant vs non-significant differential gene expression in the CovR_D53A_ mutant. Dots dispersion represents mutant-specificities. (C) Dot plots of RNA-seq fold changes for all genes in the Δ*covR* (blue) and CovR_D53A_ (red) mutants. Note the opposite trends in the total number of mild up-regulated genes (1 < log2 FC < 3) in the Δ*covR* mutant and of down-regulated genes (−3 < log2 FC < −1) in the CovR_D53A_ mutant. (D) Fold changes were plotted for each gene, organized in their order on the BM110 chromosome (x axis). The four genomic regions with an opposite transcription in the Δ*covR* (blue line) and CovR_D53A_ (red line) are highlighted with dotted boxes. (E) Validation of gene expression by qRT-PCR in BM110 strains. From the RNA-seq data, six genes were selected as induced in the two *covR* mutants (*cylJ, pil-1, fbsB, fbsA, hvgA, nucA*), induced in the Δ*covR* mutant only (*srr2*), or repressed in the two mutants (*cfb*). Means and standard deviations are calculated from three independent experiments.

The transcription of 76 and 3 genes (= 3.6% of the total number of genes) is similarly repressed or activated, respectively, in the two *covR* mutants when excluding the four prophages (Supplementary Tables S1F). Among them, genes highly repressed (FC > 10) encodes for major virulence factors such as the CC-17-specific adhesin HvgA (6), the LPxTG adhesin FbsA (32), the PI-1 pili operon and its associated regulator (22, 33), and the *cyl* operon necessary for the synthesis and export of the ß-h/c toxin (13) (Fig. 1B and Supplementary Tables S1F). In addition, CovR strongly repressed (10 < FC < 234) seven genes encoding secreted proteins (Supplementary Tables S1F), among which the FbsB adhesin (34) and the NucA endonuclease (35), as well as an overlooked set of small proteins (51 to 157 residues after signal peptide cleavage). The regulation of a large combination of cell-wall and secreted proteins establishes CovR as the central regulator of host-pathogen interaction in CC-17.

In addition to the 79 similarly regulated genes, 7 and 64 genes are differentially transcribed more than four times (|Log_2_ FC| > 2) only in the CovR_D53A_ or Δ*covR* mutants, respectively (Supplementary Tables S1). Most notably, the operon encoding for the CC-17-specific adhesin Srr2 is differentially expressed only in the Δ*covR* mutant (Fig. 1B). Quantitative RT-PCRs on independent cultures corroborate the RNA-seq fold changes in the two mutants, including the differential *srr2* transcription (Fig. 1E). The difference between the two mutants suggests indirect effects in the Δ*covR* mutant, which might be due to a crosstalk activity of CovS in the absence of its cognate regulator (36, 37), or alternatively, to the binding of the non-phosphorylated CovR_D53A_ variant on specific promoters.

### CovR binds to promoter regions along the BM110 chromosome

To identify genes directly regulated by CovR, we ectopically expressed a N-terminal epitope-tagged CovR variant (FLAG-CovR) in the BM110 Δ*covR* mutant. Expression of FLAG-CovR depends on the addition of anhydro-tetracycline (aTc) and led to a dose-dependent repression of selected genes up-regulated in the parental Δ*covR* mutant (Supplementary Fig S1). This functional FLAG-CovR variant was used for ChIP-sequencing with two independent cultures of exponentially growing bacteria in THY after induction with 50 ng/ml aTc, altogether with a similar non-induced condition (no-aTc) and an additional strain with a non-epitope tagged CovR (no-TAG) as controls.

After sequencing, analysis and manual curation, we detected 62 high-confidence reproducible peaks (mean fold enrichment FE > 4, IDR < 0.05) distributed along the 2.17 Mb BM110 chromosome (Fig 2A and Supplementary Table S2A). The summit of 42 peaks (68%) is localized between −200 and +100 bp of a start codon (Fig 2B), as expected for a transcriptional regulator. To improve the association between the binding peaks and promoters, we mapped all transcriptional start sites (TSSs) by differential RNA sequencing (dRNA-seq). In total, 1,035 TSSs were detected, including 60 associated with small or non-coding RNAs (32 intergenic and 28 antisense RNAs) and 113 TSSs inside ORFs (Supplementary Table S3). Genome-wide TSS comparison identified 953 TSSs (92%) conserved between BM110 and the reference strain NEM316 (38), increasing mapping confidence (Supplementary Table S3). CovR binding peaks were more closely associated to TSSs than to start codon (Fig. 2C), with 39 peak summits (62%) located at less than 100 bp of a TSS (Supplementary Table S2A).

**Figure 2.**
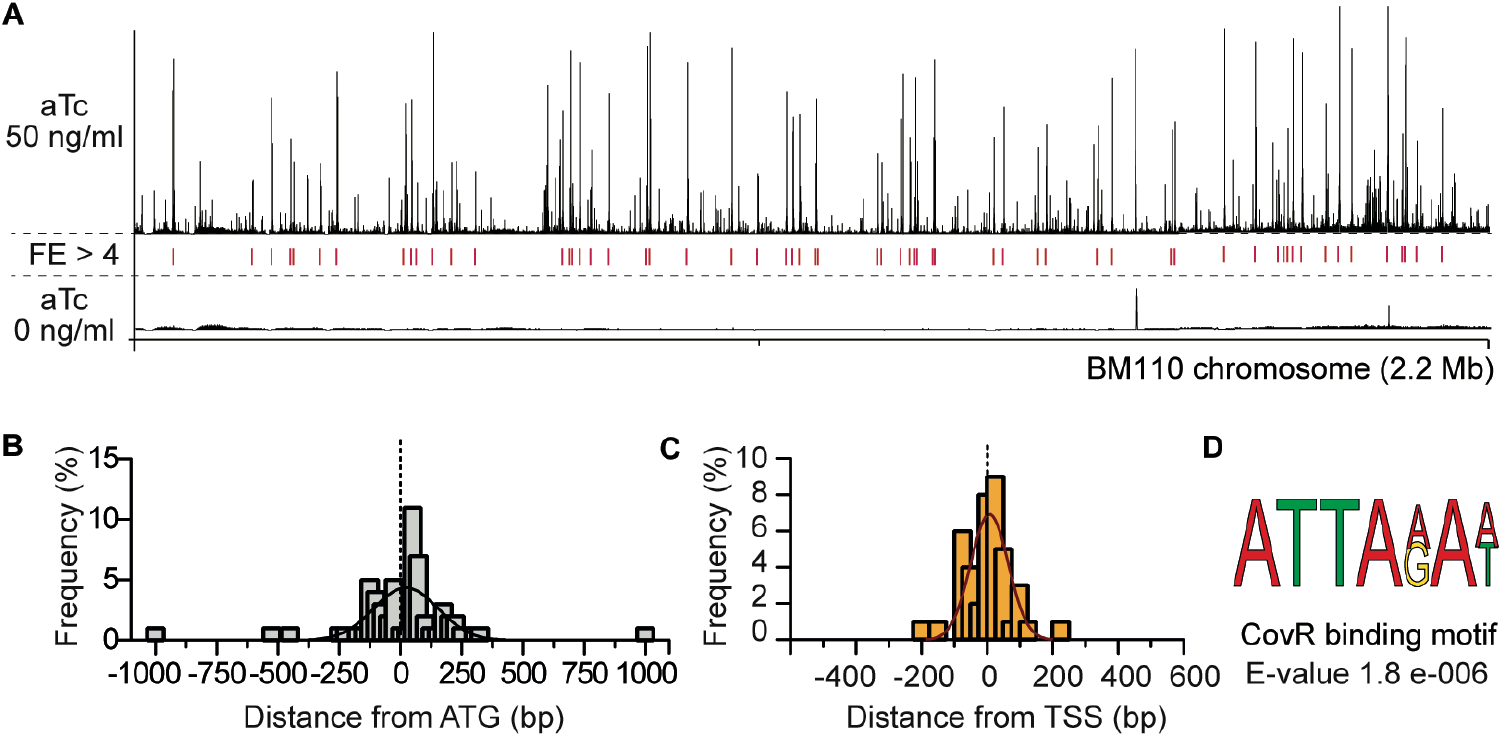
Whole-genome CovR binding on the BM110 genome. (A) ChIP-seq profile of CovR on the BM110 chromosome. Sequence reads were mapped on the chromosome after induction of the epitope-tagged FLAG-CovR with 50 (upper panel) or 0 ng/ml (bottom panel) anhydrotetracycline (aTc) in a Δ*covR* mutant. Peak height represents the mean coverage at each base pair of two independent ChIP-seq experiments. Loci with significant fold enrichment (FE > 4, IDR < 0.05) are indicated by red lines. (B) Distribution of the distance between each CovR binding peak and the nearest start codon. Distances were calculated from the summit of each CovR peak. The histogram represents the proportion of CovR binding sites (N = 62) in each sliding window of 25 bp, with an additional fitting curve. (C) Distribution of the distance between each CovR binding peak and the nearest transcriptional start site. Calculated as for (B). (D) Predicted CovR binding consensus sequence. Sequence enrichment in the 62 CovR binding loci (100 bp each) identified with the DREME software (81).

To confirm CovR binding, we selected eight promoters for *in vitro* electrophoretic mobility shift assay (EMSA). The purified recombinant protein (rCovR) binds to the eight promoters and *in vitro* phosphorylation of rCovR by acetyl-phosphate increased its affinity for all tested promoters (Fig. 3). The recombinant rCovR does not bind to the *gyrA* promoter used as a negative control and *in vitro* phosphorylation of rCovR was confirmed by Phos-Tag analysis (Supplementary Fig. S3). *In vitro* DNaseI protection assay (footprinting) on three DNA loci revealed one or two CovR-protected regions of different lengths by promoters (Supplementary Fig. S3C). Variability in number and length of CovR binding sites was previously observed (20, 23), suggesting a complex motif architecture. Indeed, a clear consensus binding motif was not detected by considering all chromosomal CovR binding loci. The most significant enriched motif is a widespread AT-rich sequence (ATTA(A/G)A(A/T)) present in 54 out of the 62 peaks, which is also the most significant motif detected by considering only peaks closely associated to a TSS (Fig. 2D).

**Figure 3.**
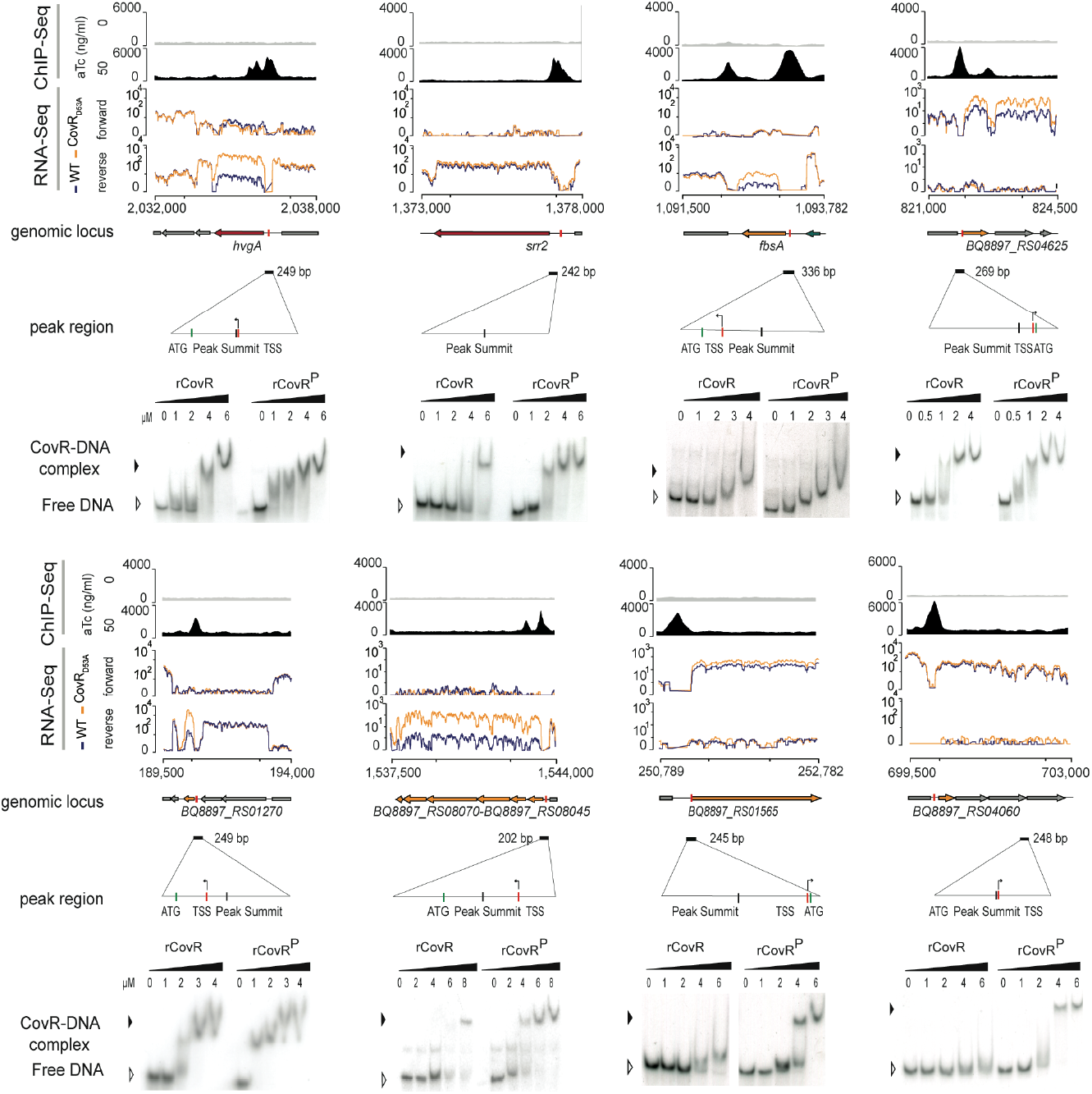
Direct and complex CovR transcriptional regulation. ChIP-seq and RNA-seq profiles showing CovR binding and the associated transcriptional signal for eight selected loci. From each locus is shown from top to bottom: *i)* the non-induced (anhydro-tetracycline (aTc) = 0 : upper gray line) and induced (aTc = 50 ng/ml : bottom dark profile) ChIP-seq profiles with the normalized sequencing coverage scale indicated on the left axis; *ii)* the normalized strand-specific RNA-seq profiles of the WT (blue) and CovR_D53A_ mutant (yellow) of the same genomic region with the chromosomal coordinates; *iii)* the schematic representation of ORFs in the locus, with the name of the regulated genes; *iv)* the position and size (202 to 336 bp) of sequences encompassing the ChIP-seq peaks, with a zoom in to highlight the position of the peak summits, the TSS, and the ATG; and *v)* the validation of CovR binding by EMSA with the recombinant purified rCovR. The formation of a rCovR-DNA complex is visualized by a delayed migration of the radiolabeled probe compared to the unbound probe (free DNA) and the affinity of rCovR to DNA is increased upon its phosphorylation by acetyl phosphate (denoted as rCovR^P^). Note that for three loci (*srr2, BQ8897_RS01565, and BQ8897_RS04060*), CovR binding is not associated to a significant transcriptional change in the CovR_D53A_ mutant.

### CovR is a direct coordinator of virulence genes expression

To characterize the direct CovR regulon, we combined the CovR binding loci identified by ChIP-seq with the transcriptional data. We chose strict criteria to define a conservative CovR direct regulon by only considering genes or operons differentially transcribed in the two *covR* mutants with a CovR binding located near the TSS (+/-100 bp) and/or the first start codon (−200 to +100 bp). This direct CovR regulon includes 21 binding loci regulating the expression of 51 genes (Table 1 and Supplementary Table S2B). All genes have an increased transcription in *covR* mutants, demonstrating the primary role of CovR as a transcriptional repressor. Notably, all but one of the highly repressed genes (N =33/34 with Log2 FC > 3) are directly regulated by CovR (Fig. 1 and Supplementary Tables S1F).

**Table 1.** The CovR direct regulon in BM110

| First gene in<br>Transcriptional<br>Units (TUs) | Number<br>of genes<br>in TUs | ChIP-seq<br>(FE) | RNA-seq<br>(mean<br>Log2 FC) | Genes<br>included in<br>TUs | Main functions |
| --- | --- | --- | --- | --- | --- |
|  |  |  |  |  | **LPxTG proteins** |
| BQ8897_RS10700 | 1 | 6,3 | 7,6 | *hvgA* | hypervirulent adhesin |
| BQ8897_RS05875 | 1 | 7,0 | 4,2 | *fbsA* | Adhesin |
| BQ8897_RS02735 | 1 | 4,2 | 1,5 | *pbsP* | Adhesin |
| BQ8897_RS03985 | 2+6 | 6,3 | 3,0 | *pil-1* operon | Adhesin and regulator (2 TU) |
| BQ8897_RS06775 | 1 | 5,5 | 1,7 | *scpB* | Serine protease (C5a peptidase) |
| BQ8897_RS02910 | 1 | 7,2 | 5,4 | *scpB2* | Serine protease (frameshifted) |
| BQ8897_RS04140 | 4 | 6,6 | 1,5 | *scpB4A* | Serine protease + Secreted small protein (134 aa) |
|  |  |  |  |  | **Secreted** |
| BQ8897_RS04080 | 12 | 4,3 | 4,6 | *cyl* operon | β-hemolysin/cytotoxin synthesis and export |
| BQ8897_RS04945 | 2 | 6,7 | 7,7 | *fbsB* | Adhesin |
| BQ8897_RS04215 | 1 | 4,4 | 3,7 | *nucA* | DNA/RNA non-specific endonuclease |
| BQ8897_RS02690 | 1 | 6,1 | 5,9 |  | Predicted esterase/lipase |
| BQ8897_RS01270 | 1 | 4,2 | 7,5 |  | Uncharacterized peptide (49 aa) |
| BQ8897_RS02625 | 1 | 6,3 | 5,0 |  | Uncharacterized small protein (141 aa) |
| BQ8897_RS09870 | 1 | 6,4 | 6,5 |  | Uncharacterized protein (586 aa - ATG miss-<br>annotation in RefSeq) |
|  |  |  |  |  | **Membrane and cytoplasmic** |
| BQ8897_RS10135 | 2 | 5,3 | 2,1 | *rgfAC* | Two-component system |
| BQ8897_RS09785 | 1 | 6,4 | 2,9 | *pepO* | Cytoplasmic endopeptidase (M13 family) |
| BQ8897_RS08070 | 6 | 5,7 | 6,7 | *pppK* | Uncharacterized - polyphosphate kinase |
| BQ8897_RS04650 | 3 | 8,1 | 2,0 |  | ABC transporter (Amino acid transport) |
| BQ8897_RS05655 | 1 | 7,1 | 3,1 |  | Membrane protein (Abx1-like) |
| BQ8897_RS04625 | 1 | 7,2 | 2,6 |  | Membrane protein (co-regulation with the |
|  |  |  |  | (*fbsC\**) | frameshifted FbsC adhesin) |
| BQ8897_RS03040 | 1 | 4,4 | 3,1 |  | Methyltransferase (frameshifted) |

At the functional level, CovR directly represses cell-wall and secreted proteins involved in GBS pathogenesis, including the HvgA, FbsA, PbsP, and Pil-1 adhesins, the C5a peptidase ScpB and related serine proteases, and the secreted NucA endonuclease and FbsB adhesin (Table 1). In addition, CovR directly represses the *cyl* operon, as well as five secreted peptides or proteins. The remaining 15 proteins of the CovR direct regulon are membrane-localized or cytoplasmic (Table 1). These proteins likely contribute to the adaptation of GBS to the host by linking virulence gene expression with metabolic uptake (amino-acid ABC transporter), proteogenic stress (*e*.*g*. polyphosphate kinase (39, 40)) and quorum-sensing (RgfAC two-component system and PepO endopeptidase (41–43)).

The direct regulon might also include the gene encoding the FbsC adhesin (Table 1). However, since the FbsC adhesin is not functional due to conserved frameshift mutations in CC-17 strains (44), we did not investigate further its co-regulation with the *BQ8897_RS04625* gene (Fig. 3). Additionally, we manually inspected the ChIP-seq profiles at the chromosomal *covR* locus. We did not observe any signal corresponding to CovR binding on its own promoter, arguing against a CovR feedback loop in BM110 (25).

### CovR signalling is embedded into an extensive network of co-regulators

Our strict definition of the CovR direct regulon encompassed 21 out of the 62 chromosomal binding sites. The remaining binding sites were sorted into a larger regulon consisting of 3 groups (Table 2 and Supplementary Table S2B). The first group includes genes differentially transcribed in one of the two *covR* mutants only, while the second group is not associated with significant transcriptional changes in *covR* mutants (Table 2). We validated CovR binding by EMSA on the operon promoters of *srr2* and of two transporters (*BQ897_RS04060* and *BQ897_RS01565*). The binding of rCovR and phosphorylated rCovR to these promoters did not differ from the binding to promoters of the direct regulon (Fig. 3). Similarly, the functions encode by groups 1 and 2 genes did not differ from the function encode by genes of the direct regulon (Table 2 and Supplementary Table S2B). The most likely explanation for the discrepancies between CovR binding and transcriptional data is a complex regulation involving additional transcriptional activators, probably overlapping (group 1) or not (group 2) the CovR binding sites. The integration of CovR signalling into a wider network of regulators is also evident by the indirect regulation (*i*.*e*. significant transcriptional changes in the *covR* mutants not associated with CovR binding) of 20 out of the 79 genes in the CovR regulon (Supplementary Table S1F).

**Table 2.**
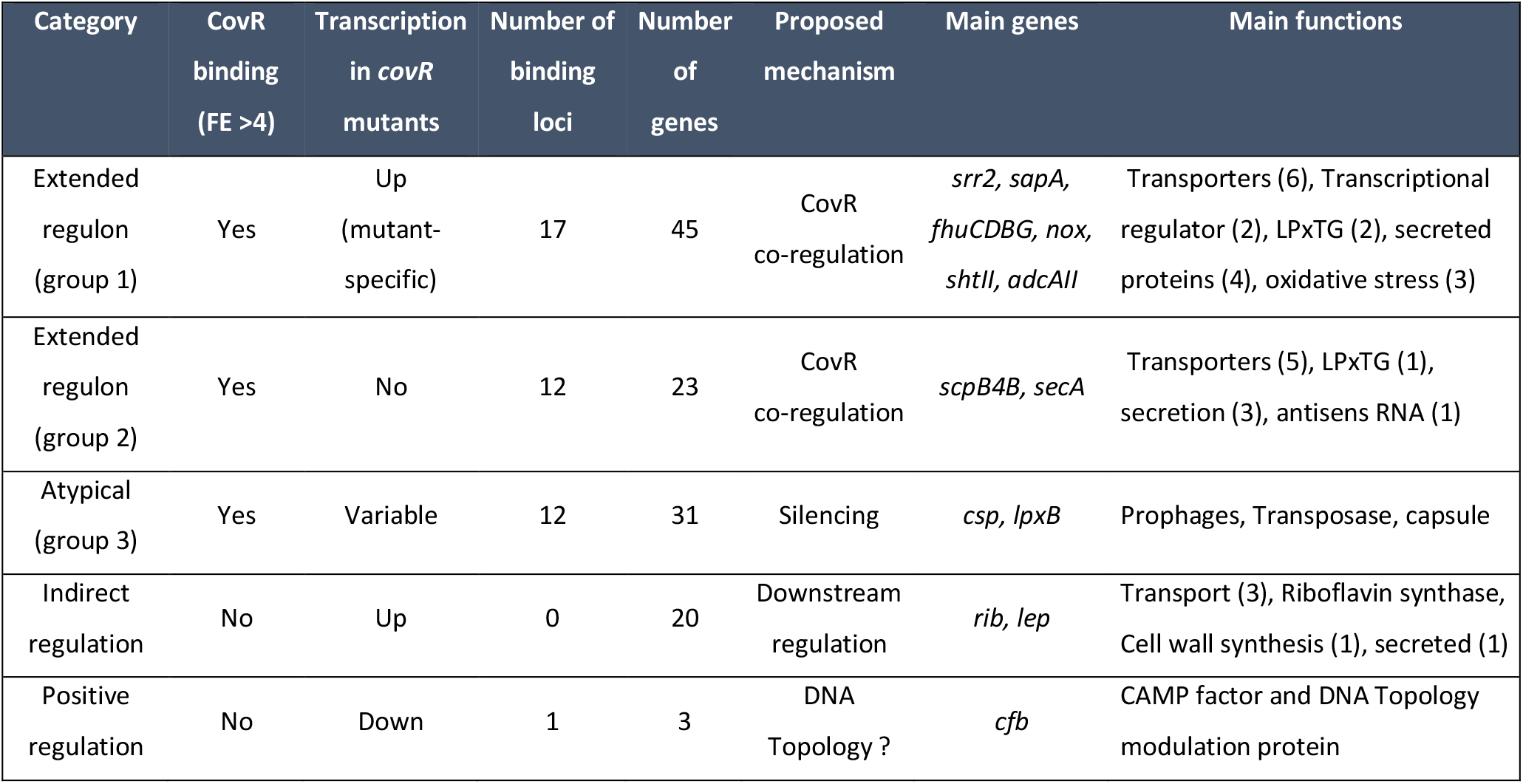
The extended CovR signaling pathway in BM110

| Category | CovR binding (FE >4) | Transcription in *covR* mutants | Number of binding loci | Number of genes | Proposed mechanism | Main genes | Main functions |
| --- | --- | --- | --- | --- | --- | --- | --- |
| Extended regulon (group 1) | Yes | Up (mutant-specific) | 17 | 45 | CovR co-regulation | *srr2, sapA, fhuCDBG, nox, shtII, adcAII* | Transporters (6), Transcriptional regulator (2), LPxTG (2), secreted proteins (4), oxidative stress (3) |
| Extended regulon (group 2) | Yes | No | 12 | 23 | CovR co-regulation | *scpB4B, secA* | Transporters (5), LPxTG (1), secretion (3), antisens RNA (1) |
| Atypical (group 3) | Yes | Variable | 12 | 31 | Silencing | *csp, lpxB* | Prophages, Transposase, capsule |
| Indirect regulation | No | Up | 0 | 20 | Downstream regulation | *rib, lep* | Transport (3), Riboflavin synthase, Cell wall synthesis (1), secreted (1) |
| Positive regulation | No | Down | 1 | 3 | DNA Topology ? | *cfb* | CAMP factor and DNA Topology modulation protein |

The third group includes the remaining 12 binding loci localized inside ORFs, in 3’ intergenic region, or associated with specific mobile elements (Table 2 and Supplementary Table S2B). Interestingly, these regions are often variable in the GBS population and several are important for pathogenesis, including the capsule operon (two CovR binding peaks in the middle and at the 3’ end of the operon), the Tn*916* element containing the *tetM* resistance gene, and the *scpB*-*lmb* locus. Notably, CovR binding is detected in prophages with atypical clusters of genes differentially transcribed in the *covR* mutants (Supplementary Fig. S2). Closer examination of these four loci showed significant CovR binding signals, either sharp (FE > 4) or diffused (1 < FE < 4), associated to a change in the transcriptional profiles of these mobile elements (Supplementary Fig. S2). This suggests that non-phosphorylated CovR contribute to the silencing of recently acquired DNA regions and that new CovR binding sites might be selected to control the expression of advantageous genes. A non-canonical mechanism of regulation might also operate for the CAMP operon, the only highly positively (Log_2_ FC < −5) CovR-regulated genes (Table 2). The promoter of the CAMP operon is the only one to display a significant ChIP-seq signal in the no-aTc control (Fig 2A), and was therefore excluded from the analysis. A second specific CovR binding signal is detected in the intergenic region and is associated to the divergently translated gene encoding a predicted nucleoid-associated protein (45), which may indicate a binding mechanism depending on DNA conformation rather than a consensus motif.

### CovR-regulated genes and promoters are under selective pressure

The binding of CovR on the promoters of the two operons encoding the HvgA and Srr2 specific adhesins directly links CovR to the hypervirulence of GBS. In addition, CovR binds to the promoter of *scpB4B*, one of the additional 68 CC-17 specific genes (4) (Supplementary Table S4). However, the *scpB4B* gene has a frameshift mutation leading to the translation of a non-functional serine protease. Interestingly, several CovR-regulated genes have frameshift mutations or internal stop codon. These mutations are localized in genes encoding proteins usually involved in GBS-host interaction such as the ScpB2 serine protease, the FbsC adhesin, a secreted endonuclease and two transporters (Supplementary Table S1F and S2A). This indicates that pseudogenization might be as important as the acquisition of new adhesins in reshaping the interaction of CC-17 with its human host.

To identify genes directly regulated by CovR and subject to selective pressures, we took advantage of the previous reconstitution of the evolutionary history of the CC-17 lineage (3, 4).

The hypervirulent lineage is a homogenous clonal complex adapted to the human host and the evolutionary pressure driving adaptation has been previously measured in the human-associated GBS population (4). In total, 24 genes associated with CovR binding accumulated more mutations than expected under a neutral model of evolution, including 15 in CC-17 specifically (Fig. 4). Additionally, a signature of positive selection with an increased frequency of non-synonymous versus synonymous mutations (dN/dS > 1) was noticeable in *srr2, scpB*, and a gene of the capsule operon (Supplementary Tables S4). A similar analysis on the intergenic regions identified 85 noncoding regions showing a significant mutational bias in the hypervirulent lineage (4). Twenty of these intergenic regions were associated with a CovR binding locus, including 9 in the CovR direct regulon (Fig. 4 and Supplementary Tables S4), potentially resulting in different CovR binding at several loci and CovR rewiring on a global scale.

**Figure 4.**
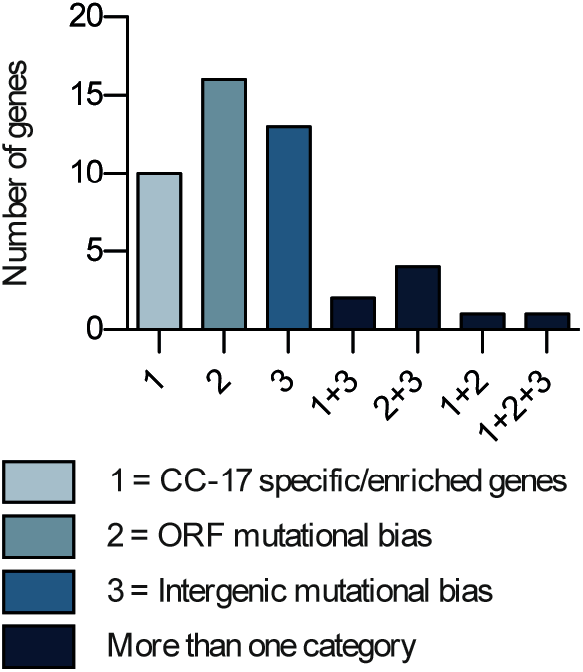
Adaptation of CovR signaling in the hypervirulent lineage. Histogram reporting the number of CovR binding loci on the BM110 chromosome associated with CC-17 specific or enriched genes (category 1), with genes showing a mutational biases indicative of adaptive evolution (category 2), and in intergenic regions with mutational biases suggestive of CovR rewiring in CC-17 (category 3). The ORFs and intergenic mutational biases in the whole GBS population have been previously calculated to reconstitute the evolution of the CC-17 hypervirulent lineage (4). Related to Supplementary Table S4.

### The plasticity of CovR signalling reshapes virulence gene expression

To quantify the specificities of CovR regulation in two strains, we did parallel RNA-seq experiments in BM110 and NEM316 (serotype III, CC-23) (46) backgrounds. The transcriptional profiles of CovR_D53A_ mutants in BM110 and NEM316 are globally similar, with a large-scale transcriptome remodelling and a distinctive set of highly activated genes (Supplementary Tables S5A and S5B). Comparative analysis on the 1,716 orthologous genes revealed distinct transcriptomes for both the WT strains and for their corresponding CovR_D53A_ mutants (Fig. 5A). Inactivation of CovR accounts for 54.4 % of the variability (PC1) between samples, while 29.8 % of the variability (PC2) is sustained by WT specificities (Fig. 5A).

**Figure 5.**
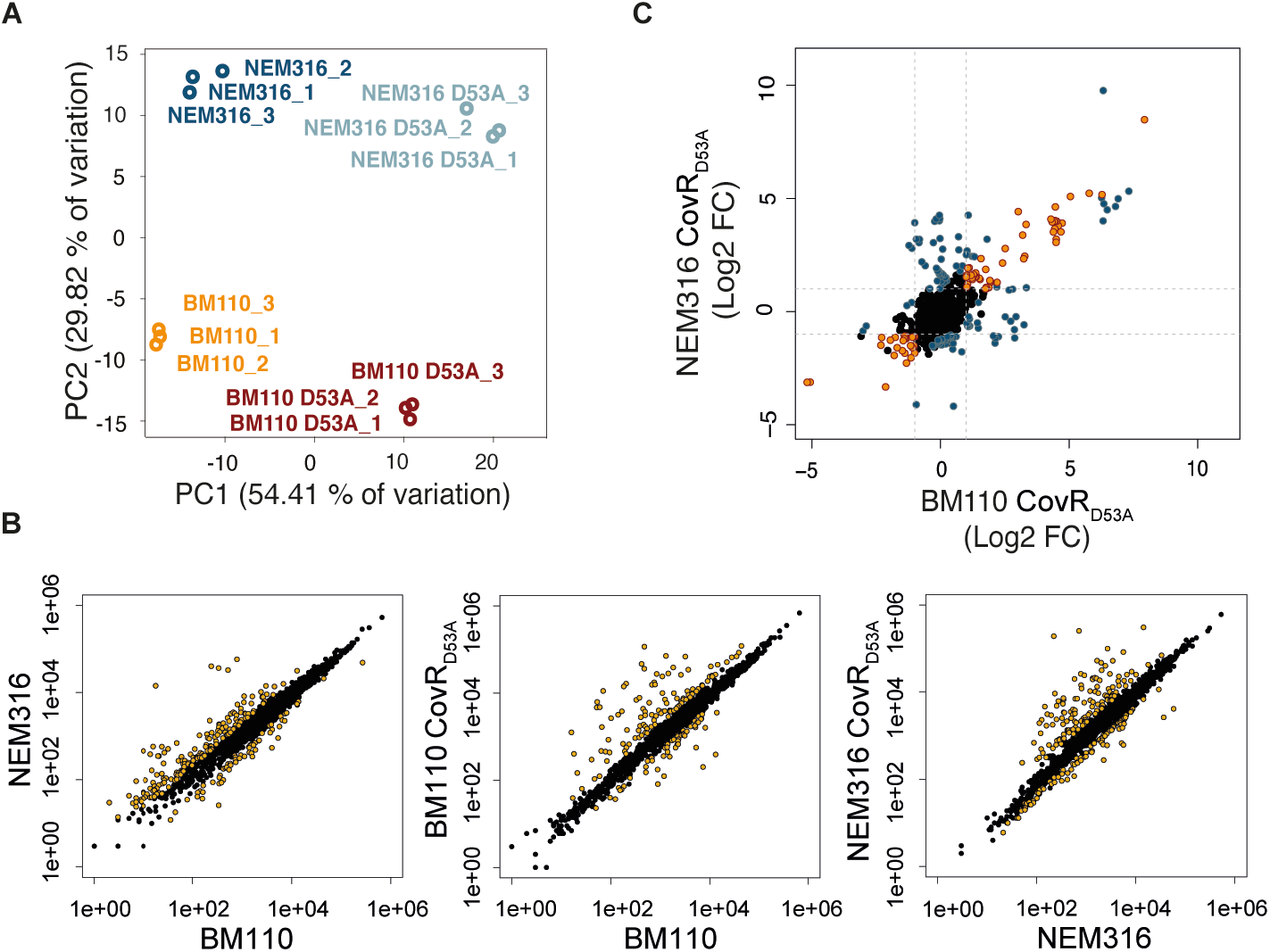
Plasticity of the CovR regulon in BM110 and NEM316. (A) Principal component analysis of RNA-seq data for the BM110 and NEM316 WT strains and their corresponding CovR_D53A_ mutants. Biological triplicates are resolved by PCA with variability sustained by CovR inactivation (PC1) and WT differences (PC2). (B) Scatter plots of the normalized read counts for each of the 1,716 homologous genes in the two WT strains, and between the WTs and their corresponding CovR_D53A_ mutants. Yellow dots represent significant differential transcription (adjusted p-value < 0.005). (C) Pairwise comparison of the fold change in the two CovR_D53A_ mutants. Genes with a similar and significant fold change upon CovR inactivation in the two backgrounds are symbolized with orange dots. Genes with a significant fold change in one mutant only or which show a significant different fold change in the two mutants are highlighted with blue dots.

The difference between the WT strains implies 172 and 60 genes with a significant increased or decreased expression (|Log_2_ FC| > 1; adjusted p-value < 0.005), respectively, in NEM316 compared to BM110 (Fig. 5B and Supplementary Tables S6A). The highest difference was observed for the expression of the direct CovR regulated gene *fbsA*, with nearly 1,000-fold induction (Log2 FC = 9.42, adjusted p-value < 10^−80^) in NEM316 compared to BM110. The *fbsA* promoter differs by 12 SNPs in the two strains, suggesting a case of CovR signalling evolution. However, *in vitro* binding of rCovR is similar on the two promoters and CovR repression is conserved in the two strains (Supplementary Fig. S4A). Therefore, the SNPs in the *fbsA* promoters do not have a direct effect on CovR binding and regulation *per se* but should be related to the gain (in NEM316) or loss (in BM110) of a binding site for an additional transcriptional activator.

The CovR_D53A_ core regulon encompasses 100 genes differentially transcribed in the same direction (activation or repression; |Log_2_ FC| > 1; adjusted p-value < 0.005) in the two strains (Fig. 5C and Supplementary Tables S6B). For 17 of these genes, the fold change associated to CovR inactivation is significantly different in the two backgrounds, and 92 additional genes are differentially expressed only in one of the two CovR_D53A_ mutants (Fig. 5C and Supplementary Tables S6C), highlighting strain-specific CovR regulation. The plasticity of the CovR regulatory pathway is especially striking for the transcription of genes encoding cell-wall anchored proteins involved in host-pathogen interaction (10). In total, we identified 27 LPxTG proteins encoding genes localized in the core or accessory BM110 genome (Supplementary Tables S7). The transcription profiles of LPxTG encoding genes in the two CovR_D53A_ backgrounds reveals a strain-specific remodelling of the bacterial surface (Supplementary Figure S5). The strain differences involve the conserved transcription of allelic variants (*bibA/hvgA*), the loss-of function mutations in highly CovR regulated genes (*scpB2* and *fbsC* in BM110, PI-1 pili operon in NEM316) and significant transcriptional difference (*pbsP, nudP cdnP, srr1/srr2 locus*) (Supplementary Figure S5).

### Strain-specificities depends on the level of CovR activation

To compare CovR binding on a genomic scale, ChIP-seq experiments in the NEM316 and BM110 backgrounds were done in parallel with two levels of CovR induction (50 and 200 ng/ml aTc) (Supplementary Table S8). In total, we detected 31 common loci associated with CovR binding in the two strains, which delineate a minimal conserved binding regulon (Fig. 6A and Supplementary Table S9A). In addition, reproducible CovR binding is observed specifically at 29 and 6 chromosomal loci in BM110 and NEM316, respectively (Supplementary Table S9A). As expected, strain-specific CovR binding occurs at the level of specific genes, such as *srr2* in BM110, and in genes localized into non-shared mobile elements. However, strain-specific CovR binding also occurs at the level of 28 loci present in the two strains. While promoters of two transporters have large deletions (68 and 73 bp in BM110 and NEM316, respectively) or SNPs, which might explain CovR binding differences, 5 binding regions are identical in the two strains on 250 bp surrounding CovR binding (Supplementary Figure S4B and Supplementary Table S9B).

**Figure 6.**
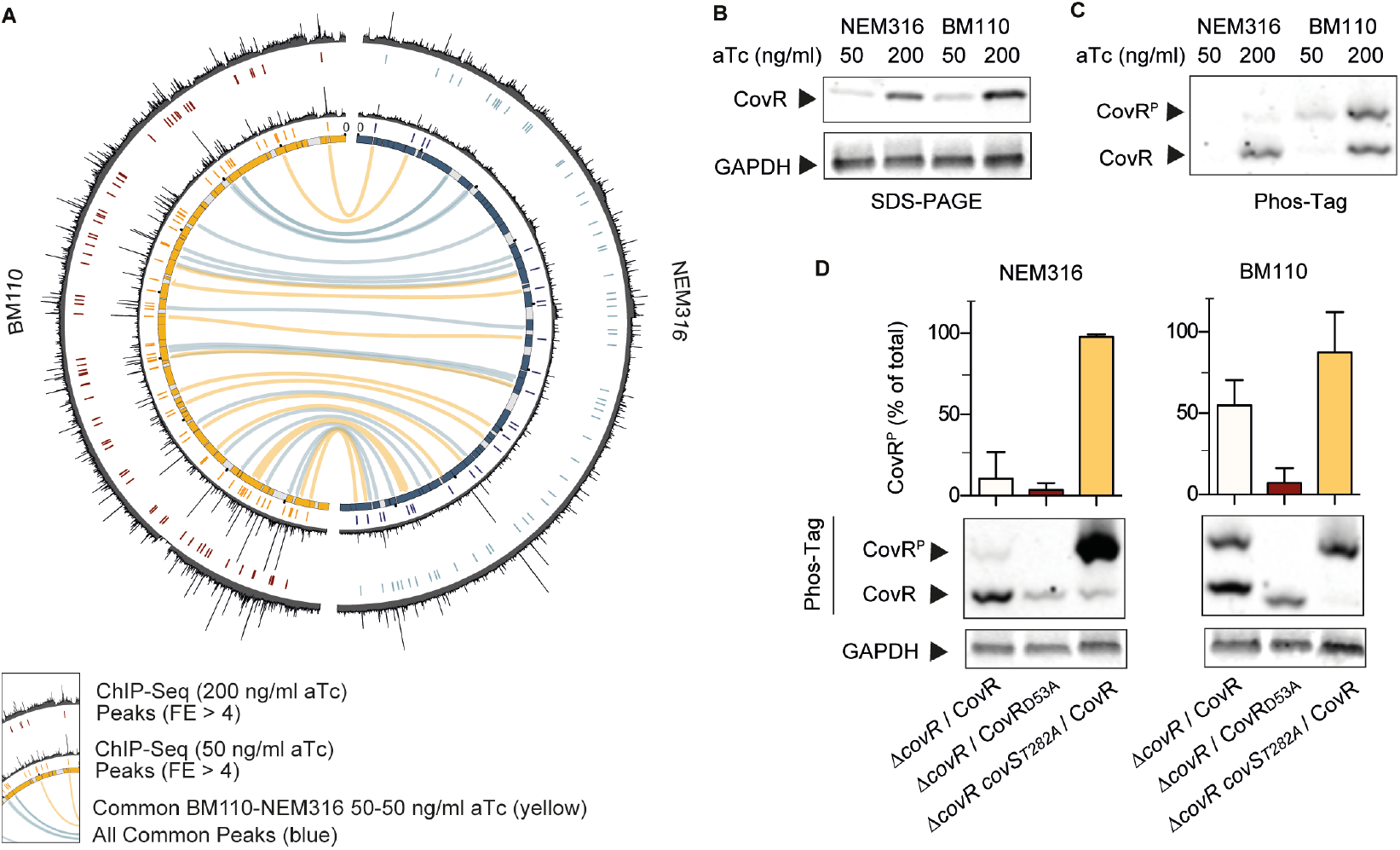
Global effect of CovR phosphorylation on chromosomal binding. (A) Shared CovR binding sites identified on the BM110 and NEM316 chromosomes. The inner circle is a symmetric representation of the BM110 (left, yellow) and NEM316 (right, blue) chromosomes with strain-specific sequences (< 90 % homology) in grey. ChIP-seq profiles for each strain are shown after induction of FLAG-CovR with 50 (inner profile) or 200 (outer profile) ng/ml anhydro-tetracycline (aTc). Significant CovR peaks (FE > 4, IDR < 0.05) are symbolized by colored traits below each ChIP-seq profile. Conserved chromosomal binding loci are represented by the inner connecting lines (in yellow: between ChIP-seq done with 50 ng/ml aTc; in blue: all ChIP-seq experiments). (B) Dose-dependent induction of FLAG-CovR by aTc in BM110 and NEM316 strains. Western blot analysis with anti-FLAG (upper panel) and anti-GAPDH (bottom panel) antibodies after SDS-PAGE electrophoresis of total protein extracts prepared from mid-exponential growing cultures in THY. (C) Level of CovR activation by phosphorylation is strain-specific. The same extracts as in (B) were analyzed with anti-FLAG antibodies after Phos-Tag electrophoresis. A delay migration denotes a phosphorylated form (CovR^P^). (D) Quantification of CovS-dependent CovR phosphorylation in NEM316 and BM110. The proportion of phosphorylated epitope-tagged CovR variants were analyzed by Phos-tag analysis after induction with 200 ng/ml aTc. The FLAG-CovR variant was used in the Δ*covR* (first rows) and Δ*covR covS*_*T282A*_ (third rows) mutants, the T282A substitution abolishing the phosphatase activity of CovS. A FLAG-CovR_D53A_ variant (second rows) unable to be phosphorylated by CovS was used in the Δ*covR* mutants. Quantification are means and standard deviation of 5 biological replicates, and a representative Phos-Tag gel with its GAPDH loading control is presented for each background.

The ChIP-seq profiles suggested a global difference in the capacity of CovR to binds chromosomal DNA between strains. This difference is most striking by comparing individual ChIP-seq experiments (Supplementary Table S9A). Analysis of ChIP-seq done with a low level of CovR induction (50 ng/ml aTc) revealed 25 significant peaks in NEM316 compared to 62 in BM110 (FE > 4, IDR < 0.05), with 15 common CovR binding regions between the two strains (Fig. 6A and Supplementary Fig. S6). By applying less stringent criteria (FE > 1, IDR < 0.05), a total of 292 and 324 significant CovR binding signals were detected in BM110 and NEM316, respectively (Supplementary Table S10), indicative of weak CovR interactions globally distributed along the chromosomes and specific CovR enrichment at regulated promoters, especially in BM110. In agreement, closer examination of the ChIP-seq profiles done with 50 ng/ml aTc revealed a lower signal to noise ratio in NEM316 compared to BM110 (Fig. 6A). In contrast, induction of CovR with 200 ng/ml aTc increases the quantity of CovR (Fig. 6B) and enhances the overall binding signal in NEM316 while it tends to increase the background signal in BM110 (Fig. 6A). In this condition, 59 and 70 CovR significant peaks (FE > 4, IDR < 0.05) were detected in NEM316 and BM110, respectively (Supplementary Fig. S6 and Supplementary Table S9A), mitigating the difference between strains.

To explain the strain difference, we hypothesized that CovR might be differentially activated by phosphorylation. Indeed, analysis of CovR phosphorylation by Phos-Tag revealed large difference between the two strains (Fig. 6C). Quantification of the different forms showed that up to 50% of CovR is phosphorylated in BM110 compared to less than 10% in NEM316 in standard growth conditions (Fig. 6D). When expressing a FLAG-CovR_D53A_ variant, only residual CovR phosphorylation is detected in the two backgrounds (Fig. 6D), probably due to the activity of the serine threonine kinase Stk1 on the CovR T_65_ residue (23). Conversely, expression of FLAG-CovR in a CovS_T282A_ mutant, in which the phosphatase activity of the histidine kinase CovS is specifically abolished (26, 27), increased CovR phosphorylation at nearly 100% in the two backgrounds (Fig. 6D). These results showed that CovS is functional in the two strains but that the basal level of CovR activation is different. This difference leads to strain-specific CovR regulation associated with a global effect on CovR binding at the genomic scale.

## DISCUSSION

Intra-species evolution of signalling pathways allows the emergence of clones associated with specific pathologies (29–31). In this study, we demonstrated that CovR is the central coordinator of GBS-host interactions by directly regulating a combination of virulence- and colonization-associated factors. In addition, our genomic analysis revealed a complex signalling pathway wired in a broader cellular network of co-regulators and the atypical regulation of genes in mobile elements. Our results also suggest that the plasticity of the CovR regulatory network underlies the ability of GBS to adapt to new environments, a condition allowing the emergence of clones associated with specific pathologies. The hypervirulence of CC-17 strains have been previously linked to the two specific adhesins HvgA and Srr2 (6, 8, 47) and we showed in this study that CovR directly regulates their expression. This was expected for HvgA since the adhesin resulted from a recombination in the central part of a gene encoding the homologous BibA adhesin present in non-CC17 clones, keeping a similar promoter with CovR binding sites (6, 20). The adhesin Srr2 is also directly regulated by CovR, but the analysis of its transcription in a mutant in which CovR is absent suggests a more complex regulation. We hypothesized that *srr2* transcription requires an activator such as the Rga-associated regulator present in the distantly related *srr1* operon of non-CC17 strains (8, 48), which might be necessary to outcompete the binding of the non-phosphorylated CovR form on the promoter. The integration of multiple regulatory proteins might be necessary to express Srr2 preferentially in the neonatal gut rather than during the invasive phase (7, 49), in contrast to HvgA which has a major role at the onset of meningitis (6).

The integration of CovR signalling with other regulators occurs at other loci than *srr2*. For instance, the PbsP adhesin encoding gene is directly activated by the SaeRS two-component system in the vagina to promote colonization (50), while *pbsP* transcription is also necessary at later stages of the infection process (51–53). Another example is the transcription of the gene encoding the FbsA adhesin (32) which depends on 3 additional regulators: RogB, RovS and Rgg (54–56). Among them, the activator RogB is present in NEM316 but absent in BM110, suggesting a genetic basis for the difference in the basal level of *fbsA* transcription between the two strains. A second level of CovR wiring is the direct regulation of transcriptional activators, such as the Ape1 activator of the PI-1 pili locus and the RgfAC two-component system. These two activators are necessary for pathogenesis but, as other CovR regulated genes, are often mutated in the GBS population, introducing variability in the network (4, 22, 41, 42).

In addition to strain differences in gene content, our high-resolution analysis revealed the important role that mobile elements have in the variability of the CovR transcriptomes (21, 25, 26). Previously, CovR has been proposed to silence a large genomic island in *Streptococcus mutans* (57). Notably, the *S. mutans* CovR is an orphan regulator which has lost its cognate histidine kinase CovS and, consequently, the CovR function is independent of its phosphorylation (57, 58). This non-canonical regulation appears conserved in GBS with clusters of prophages genes showing an opposite regulation in the Δ*covR* and CovR_D53A_ mutants. The mechanism might involve a competition between non-phosphorylated CovR and a nucleoid-associated protein (NAP), as observed in *S. mutans* (57, 58), and might also be co-opted to regulate other loci such as the capsule operon which shows strain-specific CovR regulation (59). The regulation of mobile elements by NAPs or by the recruitment of an ancestral regulatory network, such as PhoP/Q or OmpR/EnvZ two-component systems, have been extensively described in Gram-negative bacteria (28, 60). Our results reveal that CovR is involved in the regulation of genes in mobile genetic elements in GBS, but the mechanism as a silencer or anti-silencer need to be characterized (61).

Even if the non-phosphorylated CovR form seems to exert a regulatory role *in vivo* at specific loci, CovR phosphorylation increases its affinity for DNA *in vitro* leading to *in vivo* CovR enrichment on specific promoters, as expected for a canonical two-component system (62). The global regulators of the PhoP/Q family binds to complex promoters with binding sites that can vary in number, sequence, location and orientation (63, 64). This variability ensures a dynamic expression of the regulated genes, with promoters having the highest affinity for the regulator being the first to be activated or repressed (63). The unanticipated difference in CovR phosphorylation state between the two GBS strains offers a glimpse into this temporal hierarchy of regulated genes. The CovR binding regulon in BM110 is likely close to be exhaustive while in NEM316 only promoters with the highest affinities for a phosphorylated CovR might have been identified (63). An interesting instance is the *cyl* operon encoding the ß-h/c toxin which is directly regulated by CovR (20, 21, 65). *In vivo* binding of CovR is highly significant in the BM110 strain, but is below the threshold (FE > 4) in the NEM316 strain. This correlates with the lower haemolytic activity and the higher level of CovR phosphorylation in BM110 compared to NEM316. The paradox of a more efficient repression of virulence genes in the hypervirulent clone might in fact reflect the clinical characteristic of CC-17 strains which are associated with a delayed colonization of the neonatal gut rather than an increase pathogenicity caused by toxin expression (7, 49, 66).

The difference in the basal level of CovR phosphorylation has likely an effect on the dynamics of the response to external stimuli, for instance in the acidic phagolysosome (67, 68). Sensing and CovR activation is mediated by CovS and two additional proteins, the serine-threonine kinase Stk1 (23) and the CovS-interacting protein Abx1 (26). While Stk1 and Abx1 are identical in the NEM316 and BM110 strains, CovS differs by one residue between the two strains. We cannot exclude at this stage that the CovS polymorphism, a valine to alanine substitution at position 112 localized into the extracellular loop, is the cause of the differential CovR phosphorylation in the two strains. An alternative hypothesis will be that CovR phosphorylation depends on an additional, but yet uncharacterized, regulator. Deciphering the mechanism of CovR activation and its variation in the GBS population is therefore essential to accurately compare the evolution of the CovR regulatory pathway.

Regulatory evolution and network plasticity are two fundamental mechanisms allowing to generate phenotypic heterogeneity at the species level. The rewiring of a common regulator to control specific genes necessary in a specific environment (29) or mutations in global regulators to reshape the entire network (30) allow the emergence of clones associated with particular diseases. In Group A *Streptococcus* (GAS), CovR/S polymorphisms are a major determinant of the phenotypic heterogeneity in the population (31, 69) and mutations in the associated regulator RocA are associated with the unusual severity of invasive infections by M3 serotype (70–72). At the individual level, *covR/S* loss of function mutations have been also frequently identified in GAS clinical isolates (19, 30, 73), but these mutations have a cost and decrease host-to-host transmission of the hyper-invasive isolates (30, 74). Similarly, hyper-invasive *covR/S* null mutants have been occasionally identified in GBS, especially in strains causing *in utero* infections (13), and the CovR transcriptomes are markedly different in strains of different clonal complexes (21, 25, 26). Here, we have demonstrated that CovR is the direct and global regulator of host-bacteria interaction. Due to its central function, the whole CovR network is subject to selective pressure and has evolved in the streptococcal population to generate diversity, ultimately resulting in the selection of clones with strain-specific CovR regulation associated with specific infections.

## MATERIALS AND METHODS

### Strains and growth conditions

BM110 (Serotype III, CC-17) and NEM316 (Serotype III, CC-23) strains are representative of two of the five main GBS clades associated with human infections (3, 46). All strains and plasmids used in this study are detailed in the Supplementary Table S11, and standard growth conditions are defined as cultures in Todd-Hewitt Yeast (THY) buffered with 50 mM Hepes (pH 7.4) incubated at 37°C in static condition. Columbia agar supplemented with 10% horse blood and Granada medium (BioMerieux) were used for propagation and for visualisation of ß-hemolytic activity and pigmentation, respectively. Erythromycin and kanamycin (Sigma-Aldrich) are used for plasmid selection and maintenance at 10 and 500 µg/ml, respectively, while anhydrotetracycline (Sigma-Aldrich) is used for conditional expression. For *Escherichia coli*, LB medium were used with ticarcillin (100 µg/ml), chloramphenicol (30 µg/ml), erythromycin (150 µg/ml), or kanamycin (25 µg/ml) when appropriated.

### Plasmids and strains construction

Oligonucleotides and plasmids construction are detailed in the Supplementary Table S11. Briefly, for the epitope-tagged expression vector, a synthetic DNA containing a translational initiation site, a start codon, the 3xFLAG epitope, a flexible linker, three stop codons in the three reading phases and a transcriptional terminator was synthesized and cloned into a pEX vector (MWG Genomics). An inverse PCR on the pEX vector and a PCR on genomic DNA, followed by Gibson assembly, were used to precisely clone the *covR* sequence between the linker and the stop codons. The *covR*-synthetic DNA fragment was excised from the pEX vector by BamHI digestion and cloned into the aTc inducible expression vector pTCV_P_tetO_ (44). The construction of the Δ*covR* mutants were previously reported in NEM316 (26) and BM110 (53), as well the construction of the NEM316 CovR_D53A_, and CovS_T282A_ mutants obtained by precise chromosomal substitutions (26). The BM110 point mutants were constructed as described in NEM316 with the pG1 shuttle thermosensitive plasmid (26). For recombinant rCovR purification, the *covR* sequence was cloned into the pET-24a expression vector (Life Technologies) and transformed into BL-5 *E. coli* strain for expression.

### RNA- and dRNA-sequencing

RNAs purification for RNA-seq were done from three independent cultures, with replica done in different days, in 10 ml of THY, 50 mM Hepes pH 7.4. RNA stabilization reagents (RNAprotect, Qiagen) were added at mid-exponential phase (OD_600_ 0,5-0.6) for 5 min at ambient temperature. Cells were harvested at 4°C, washed with 1 ml cold PBS, and the bacterial pellets stored at minus 80°C. Cells were mechanically lysed and total RNA extracted following manufacturer instructions (FastPreps and FastRNA ProBlue, MP Biomedicals). Residual DNA were digested (TURBO DNase, Ambion) and samples qualities were validated (Agilent Bioanalyzer 2100, Qubit 3.0, Life Technologies) before rRNA depletion, libraries construction and sequencing (Ribozero rRNA, TruSeq Stranded mRNA, Hiseq2500, Illumina). RNA purification for dRNA-seq to determine TSS positions in the BM110 strain were done, processed and analysed exactly as described in the NEM316 strain (38). The specificities of the dRNA-seq protocol are a Tobacco Acid Pyrophosphatase (TAP) treatment of RNAs and the use of a specific 5′ adapter to differentiated primary transcripts and processed RNAs. For qPCR analysis, RNAs were prepared as for RNA-seq. Standard reverse transcription and quantitative PCR were done as described (Biorad) (26).

For RNA-seq, single-end strand-specific 65 bp reads were cleaned (cutadapt version 1.11) and only sequences at least 25 nt in length were considered for further analysis. Alignment on the corresponding reference genomes (Bowtie v1.2.2 with BM110 RefSeq NZ_LT714196 and NEM136 RefSeq NC_004368) (75) and gene counts data (featureCounts, v1.4.6-p3, Subreads package; parameters: -t gene -g Name -s 2) were analysed with R (v3.6.1) and the Bioconductor package DESeq2 (v1.26.0) (76). Normalization and dispersion were estimated and statistical tests for differential expression were performed applying the independent filtering algorithm. A generalized linear model including the replicate effect as blocking factor was set in order to test for the differential expression between the mutant and the WT strains. For each comparison, raw p-values were adjusted for multiple testing according to the Benjamini and Hochberg procedure (77) and genes with an adjusted p-value lower than 0.005 were considered differentially expressed. The coverage profiles were obtained for each strand using bedtools (v2.25.0), normalized using the DESeq2 size factors and then averaged across the biological replicates.

### Chromatin immunoprecipitation and sequencing

Cultures of the BM110 and NEM316 Δ*covR* mutants containing the pTCV-P_tetO_-FLAG-*covR* or the pTCV-P_tetO_-*covR* vector were done in parallel with independent duplicates for each condition. Overnight cultures in THY, Hepes 50 mM, kanamycin 500 µg/ml, were inoculated (1/50) in 100 ml of fresh media supplemented with the indicated concentration of aTc, and incubated until mid-exponential growth phase (OD_600_ 0,5-0,6). Crosslinking were done by the addition of 1% formaldehyde for 20 min at room temperature under agitation, followed by quenching with 0,5 M glycine for 15 min. Bacteria were harvested, washed two times in ice-cold Tris-Buffered saline (20 mM Tris/HCl pH 7.5, 150 mM NaCl) and resuspended in 1 ml Tris-Buffered saline supplemented with protease inhibitor (cOmplete Protease Inhibitor, Roche). Bacterial lysis was done by enzymatic cell wall degradation (30 min at 37°C with 10K/ml mutanolysine, Sigma) followed by mechanical disruption (FastPrep-24, MP biotechnological) at 4°C with 0,1 mm glass beads (Scientific Industries, Inc). After centrifugation (4°C, 5 min), 500 µl of supernatants were diluted with 500 µl of cold immunoprecipitation buffer (50mM HEPES-KOH, pH 7.5, 150mM NaCl, 1mM EDTA, 1% Triton X-100, 1mM fresh PMSF) and the chromatin was fragmented by sonication (Covaris S220) for 20 min in milliTUBE (1ml with AFA Fiber). Aliquots of 100 µl were collected to measure DNA fragmentation and to confirm CovR expression by agarose gel electrophoresis and Western blot with anti-FLAG antibodies, respectively.

Chromatin immunoprecipitation were done with 40 µl of Anti-FLAG M2 magnetic beads (Millipore) for 2h at 4°C under constant agitation. Beads were successively washed with four buffers (twice in immunoprecipitation buffer, twice in 50mM HEPES-KOH pH 7.5, 500mM NaCl, 1mM EDTA, 1% Triton X-100, 1mM fresh PMSF, once in 10mM Tris-HCl pH 7.5, 250mM LiCl, 1mM EDTA, 0.5% NP-40, 1mM fresh PMSF, and once in TE buffer 10mM Tris pH 7.5, 1mM EDTA). CovR elution were done in 50mM Tris-HCl pH 7.5, 1mM EDTA, 1% SDS pH 8.0 and confirmed with aliquots analysed by Western blot. Samples were treated with RNAse (Sigma) for 30 min at 37°C and reverse crosslinking was carried out by an overnight incubation at 65°C with 0,1 mg/ml proteinase K (Eurobio). Magnetic beads were discarded and DNA purified (QIAPREP, Qiagen) and quantified (Qubit 3.0, Invitrogen). DNA libraries preparation with 16 cycles of PCR amplification and sequencing were done following manufacturer instructions (TruSeq ChIP-Library kit, NextSeq 500/550, Illumina).

Quality controls, trimming and genome mapping of single end sequencing reads (75-bp) were first proceeded similarly to the RNA-seq reads. A step was applied to filter duplicated reads (Picard-tools v2.8.1, Samtools, v1.6) and a strands cross-correlation metrics step (phantompeakqualtools, R, v3.0.1) was applied for quality metrics and to evaluate fragment length before peak calling (Macs v2.1) (78, 79). The corresponding no-tag samples were used as controls and only peaks with a p-value inferior to 0.01 were considered. Reproducible peaks between independent replicates were identified with an expected rate of irreproducibility discovery threshold of 0.05 (IDR, v2.0.2) (80). Functional assignation of peak summits to ATG and TSS were done (BEDtools v2.25.0) before manual validation on normalized read coverage generated with custom bash scripts and visualized with Integrated Genome Viewer (v2.3.25) and Geneious (Biomatters Ltd, v2019.2.3). DREME (81) was used to find enriched motifs using 100 bp of sequences centred on peak summits.

### Recombinant rCovR purification

The recombinant rCovR with a C-terminal histidine tag was purified from 800 ml of culture of Bli5 *E. coli* containing the corresponding pET-24a-*covR* expression vector by column chromatography. Induction was done on growing cultures at OD_600_ = 0.8 by the addition of IPTG (0.1 mM) followed by an overnight incubation at 20°C. Bacterial cells were collected by centrifugation, frozen at −20°C, resuspended in 40 ml of 50 mM Na2HPO4/NaH2PO4, 300 mM NaCl, pH 7.0, and lysed through two passages on a cell disruptor. After centrifugation, supernatants were filtered (0,22 µM Steriflip, Merck) and incubated for 20 min under constant rotation with 3,5 ml of pre-washed Ni-NTA superflow beads (Qiagen). Beads were collected by centrifugation, washed two times for 10 min with 20 ml of fresh buffer, and elution of rCovR from beads were done on the top of a size exclusion column (Biorad) with 50 mM Na2HPO4/NaH2PO4, 300 mM NaCl, 150 mM imidazole, pH 7.0. Fractions containing rCovR were pooled and desalted (PD-10 columns, GE Healthcare) with 50 mM Tris-HCl, 500 mM NaCl, pH 8. Aliquots of rCovR were conserved at −20°C in buffer supplemented with 30% glycerol, pH 8.

### CovR phosphorylation and *in vitro* binding

For *in vitro* rCovR phosphorylation, up to 5 µg of rCovR was incubated for 60 min at 37°C with 20 mM MgCl2 and 35 mM lithium salt acetyl-phosphate (Sigma), and phosphorylation was confirmed by Western analyses with anti-His tag antibodies after electrophoresis in 12.5 % Phos-Tag SDS polyacrylamide gels (SuperSep Phos-Tag, Wako Pure Chemical Industries Ltd) for 2 hours (100V, 30 mA) in Tris-glycine buffer on ice. The negative control is the heated (100°C, 1 min) sample, removing the phospho-labile phosphoryl group. *In vivo* CovR phosphorylation were similarly analysed with rabbit anti-FLAG antibodies (1:1000) after electrophoresis of 20 µg of total GBS protein. Mouse anti-GAPDH antibodies were used as loading controls. Fluorescent secondary antibodies (goat anti-rabbit and anti-mouse IRDye 800 CW, Licor Biosciences) were used and signals revelation and quantification were done with Odyssey Imaging system (Licor Biosciences) on at least five independent protein extracts. Electrophoretic mobility shift assay (EMSA) were done with PCR probes produced with a forward primer previously radiolabelled with [γ-32P]-dATP by the T4 polynucleotide kinase (New England Biolabs). Protein-DNA interaction was performed with variable concentrations of rCovR, radiolabelled probe, 0.1 µg/µl of Poly(dI-dC) (Pharmacia), and 0.02 µg/µl BSA in binding buffer (25 mM Na2HPO4/NaH2PO4 pH 8, 50 mM NaCl, 2 mM MgCl2, 1 mM DTT, 10% glycerol) for 30 min at room temperature. Samples were separated onto a 5% TBE-polyacrylamide gel for 1 h 30 min and revealed by autoradiography. For each probe, EMSA were done with the same aliquot of rCovR without and with extemporaneously phosphorylation. DNase I protection assays (footprinting) were done in similar condition and as previously described (20).

## Data availability

All sequencing data (RNA-seq, dRNA-seq, ChIP-seq) have been submit to the repertory GEO (with private access until publication).

## Author contributions

MVM, IRC, OS, MG, PAK, and AF performed experiments, MVM, MD, HV, IRC, RL, PG, CC, and AF analyzed data, PTC and AF conceived and designed the study, MVM, PTC and AF wrote the paper with contribution of all co-authors.

## Acknowledgements

We thank R. Koszul, M. Marbouty and A. Thierry for their advices on DNA sharing, sequencing and analysis.

