## Supplementary figures for "The CovR regulatory network drives the evolution of Group B *Streptococcus* virulence"

**Supplementary Figure S1. Complementation of the *covR* mutant phenotypes by the inducible epitope-tagged FLAG-CovR variant.**

**Supplementary Figure S1**

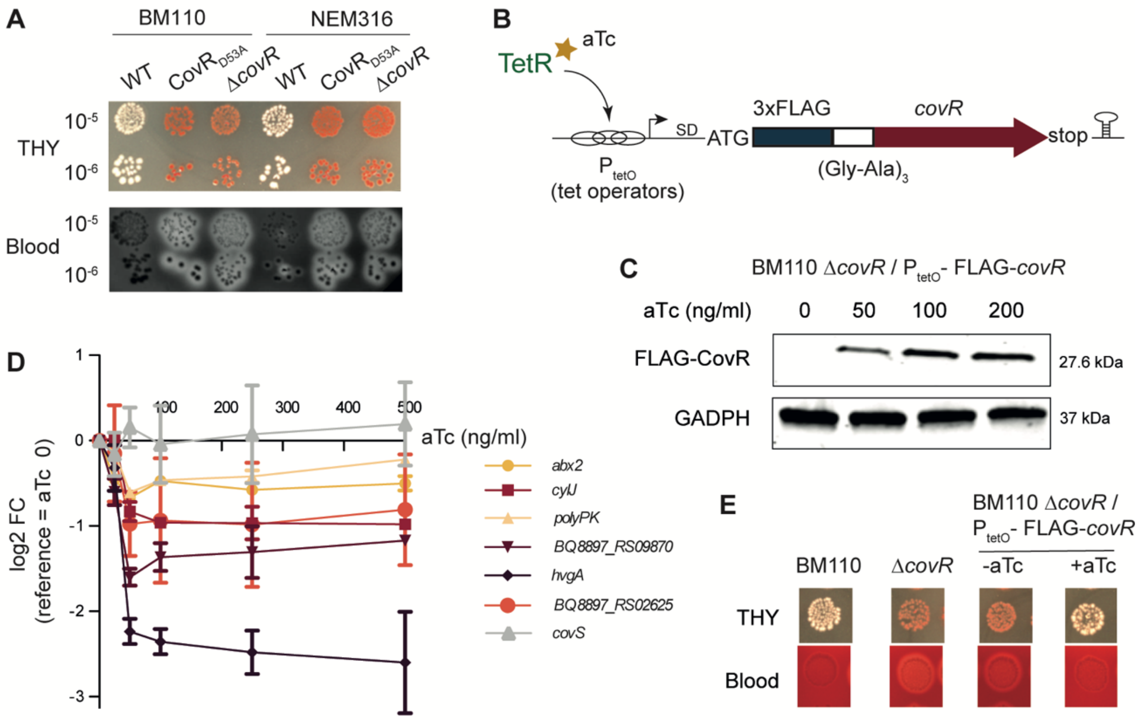


(A) Pigmentation and ß-hemolytic phenotypes of *covR* mutants. Dilutions of overnight cultures ($10^{-5}$ and $10^{-6}$) of the BM110 (CC-17) and NEM316 (CC-23) wild-type strains and of their corresponding Δ*covR* deletion and CovR$_{D53A}$ substitution mutants were spotted on rich media (THY) and Columbia agar supplemented with horse blood (Blood). Photos were taken after 24–36 h of growth at 37°C.

(B) Schematic representation of the epitope-tagged FLAG-CovR system. The epitope-tag (3xFLAG) is linked in frame to the 5' end of the *covR* sequence by a flexible linker (Gly-Ala x 3). Transcription is dependent upon the binding of anhydro-tetracycline (aTc) to the trans-activator TetR, leading to the binding of TetR to the P$_{tetO}$ promoter containing tet operators sequences upstream the transcriptional start site. The *tetR* gene is cloned in the same pTCV-$P_{tetO}$ vector (not represented) as the $P_{tetO}$-FLAG-*covR* cassette.

(C) Dose-dependent induction of FLAG-CovR by aTc. Western analysis of total protein extracts with anti-FLAG and anti-GAPDH antibodies after cultures of the BM110 Δ*covR* mutant containing the pTCV-tetO-FLAG-*covR* expression vector with increasing concentration of aTc.

(D) Repression of negatively CovR-regulated genes by the epitope-tagged FLAG-CovR variant. Induction of FLAG-CovR in the Δ*covR* / pTCV-$P_{tetO}$-FLAG-*covR* strain was done with increasing concentration of aTc (0, 25, 50, 100, 250, 500 ng/ml) and RNA were prepared from mid-exponentially growing cultures. RT-qPCR were done on selected genes up-regulated in the parental Δ*covR* mutant and on the *covS* gene as a control. Results are reported as the fold change difference (log2 FC) between the induced and the non-induced (aTc = 0) condition, with *gyrA* as the reference gene. Mean and standard deviation are calculated from three biological replicates.

(E) Phenotypic complementation of the pigmentation and ß-hemolytic phenotypes of the BM110 Δ*covR* mutant. The BM110 Δ*covR* / pTCV-$P_{tetO}$-FLAG-*covR* strain was spotted as in (A) on THY and Columbia Blood Agar without (-aTc) or with 500 ng/ml aTc supplementation (+aTc).

**Supplementary Figure S2. Atypical CovR-regulation of genes in mobile elements.**

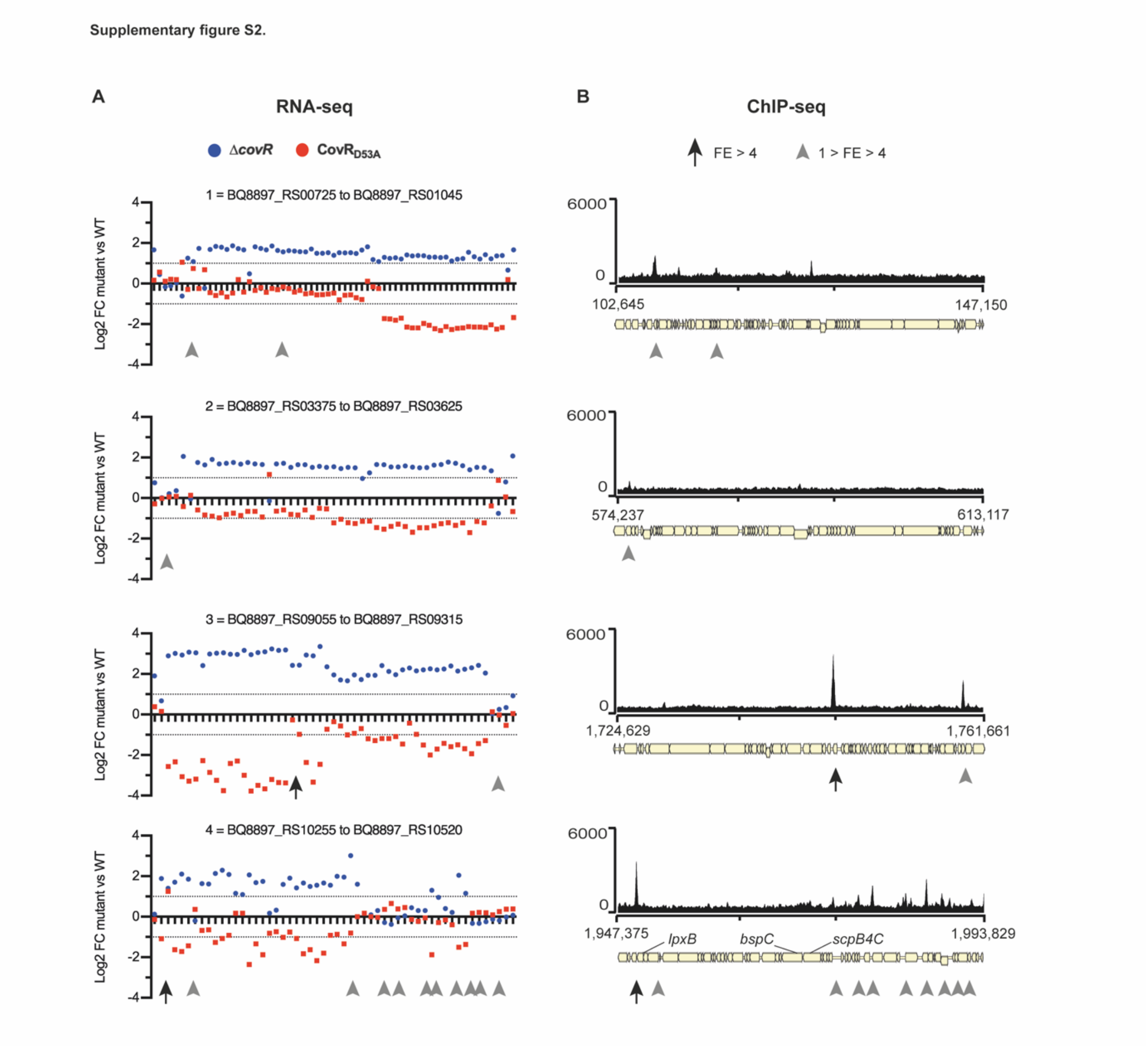


(A) Zoom in the four genomic regions highlighted in Fig. 1D with genes IDs.

(B) Corresponding ChIP-seq profiles obtained after induction of FLAG-CovR with 50 ng/ml in a BM110 Δ*covR* background. Significant (IDR < 0.05) CovR peaks are depicted with black arrows (FE > 4) or grey arrowheads (1 < FE < 4) below the schematic ORF representation of the loci. The CovR binding sites are reported in (A) for comparison with RNA-seq data.

**Supplementary Figure S3.** ***In vitro*** **phosphorylation and binding of rCovR.**

**Supplementary Figure S3.**

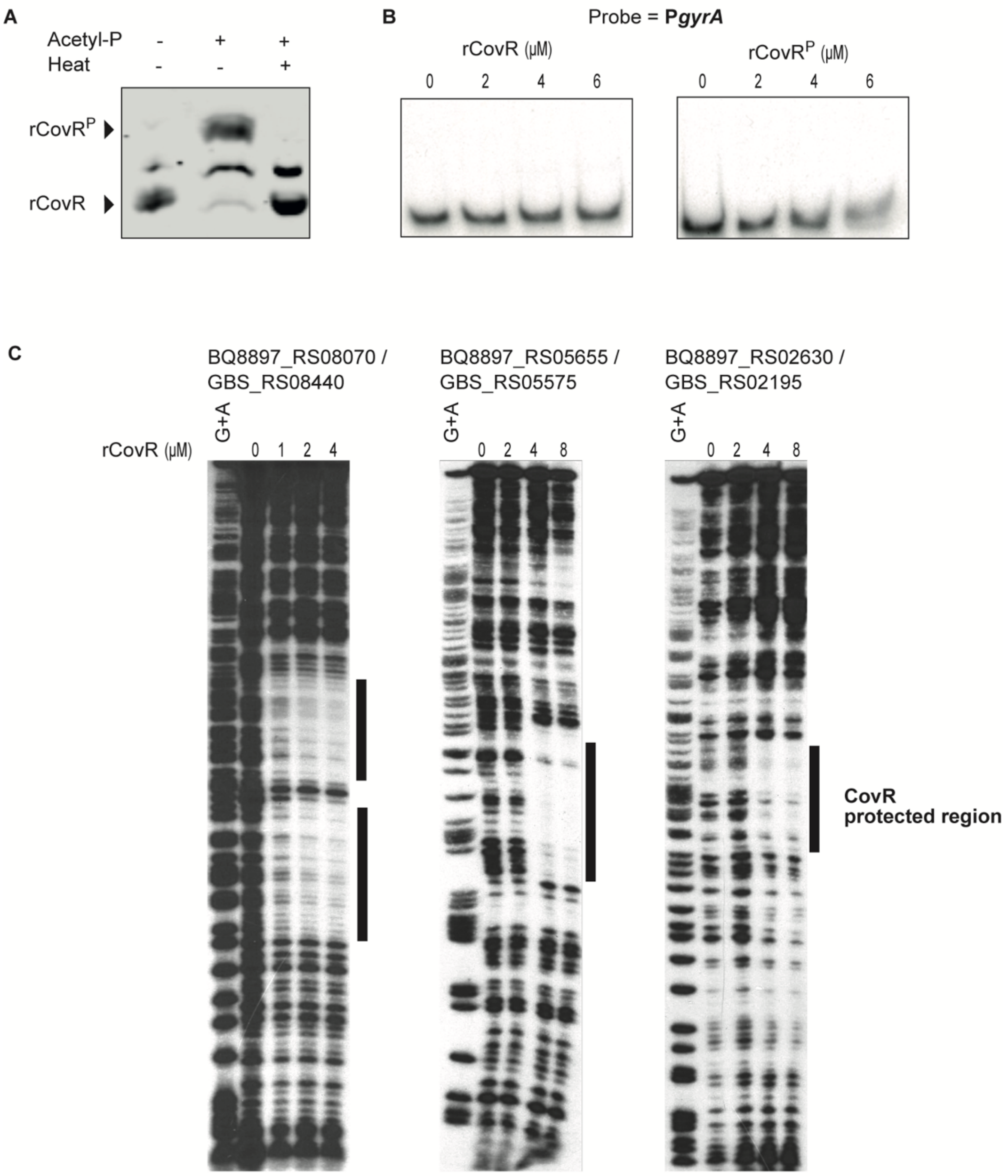


BQ8897_RS08070 / GBS_RS08440 — +1 — ATATATTTATTTTTGCTTAATTTATTGTTAATATttatATATAAATTAATATTTAATTAAACA

BQ8897_RS05655 / GBS_RS05575 — +1 — GTATAAAGTTAGATAAAAGTGAGATATTTATGAGA(G/A)

BQ8897_RS02630 / GBS_RS02195 — TATAATGATTGACATTTTTATCTAAT(T/G)AA...(17bp) +1

(A) Control of rCovR phosphorylation by acetyl phosphate. Recombinant rCovR was incubated without (-) or with (+) acetyl-phosphate for 1 hour at 37°C. One-half of the sample incubated with acetyl phosphate was subsequently heated at 100°C for 1 minute to remove the labile aspartate phosphorylation. Samples were separated on a 12% Phos-Tag SDS polyacrylamide gel and transferred on membrane before revelation with anti-His antibodies. The Phos-tag gel delays the migration of phosphorylated proteins compared to non-phosphorylated proteins.

(B) Negative binding control of rCovR on the $P_{gyrA}$ promoter by EMSA. Increased quantity of purified recombinant protein without (rCovR) or with ($rCovR^P$) phosphorylation by acetyl phosphate are mixed with radiolabeled $P_{gyrA}$ probe and separated on 5% PAA-gel before autoradiography revelation. Related to Figure 3.

(C) DNase I protection assay with rCovR on three genomic loci. Single-end radiolabeled probe are mixed with increasing quantity of rCovR before DNaseI treatment. Binding of rCovR on DNA is visualized by the DNA sequences protected from DNase I degradation ('footprints' highlighted by vertical black lines) after separation on Maxam-Gilbert sequencing gels. For the 3 loci, the corresponding rCovR protected sequences are given with the gene IDs in BM110 (BQ8897_RSxxxxx) and NEM316 (GBS_RSxxxxx). The TSSs are indicated by an arrow (+1), the ATG start codon by a box when it is within the sequence, and the SNPs between BM110 and NEM316 by the nucleotides in brackets. Note that different rCovR batches have been used for the three experiments, hampering a direct comparison of rCovR affinities between promoters.

**Supplementary Figure S4. Promoter mutations in direct CovR regulated genes.**

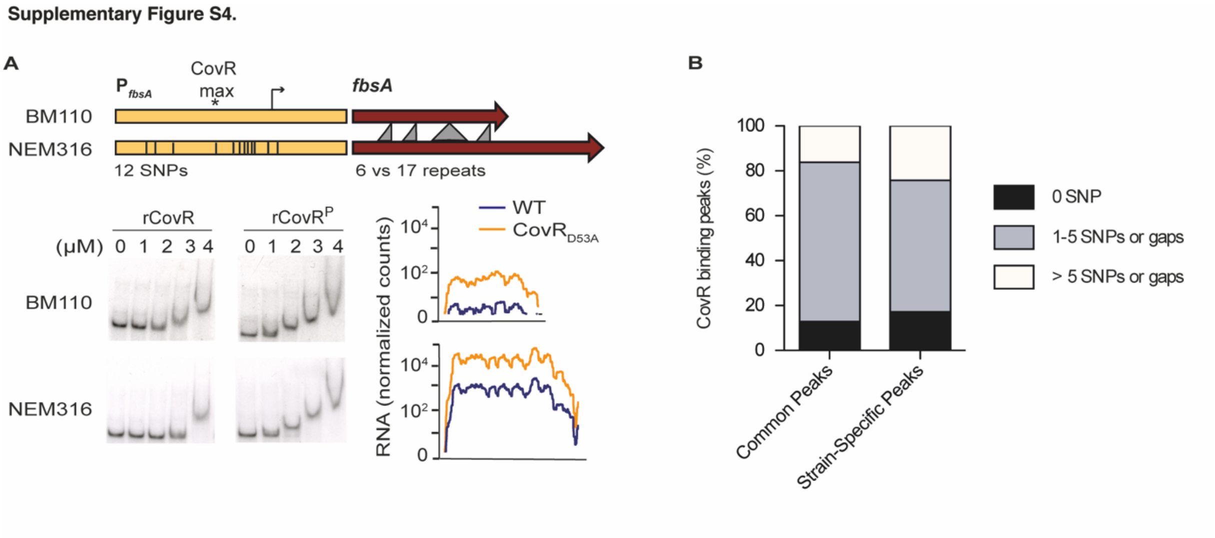


(A) CovR regulation of the FbsA adhesin encoding gene in BM110 (CC-17) and NEM316 (CC-23) strains. Schematic representation of the *fbsA* gene (dark red) and promoter (yellow) in the two strains showing the gene length difference leading to the translation of a protein with a different number of a repeated motif, and the 12 SNPs in the promoter region (336bp). Bottom: binding of non-phosphorylated and phosphorylated rCovR to the two promoters by EMSA and normalized RNA-seq data for the two WT strains and their corresponding $CovR_{D53A}$ mutants. Note that the two strains differ by their basal level of *fbsA* transcription and not by CovR binding or regulation.

(B) Cumulative percentage of CovR binding loci (250 bp centered on the maximum ChIP-seq signal) with 0 (black), up to five (grey) or more than five SNPs or gaps (white) between BM110 and NEM316 sequences. Left: CovR binding loci detected in BM110 and NEM316; Right : CovR binding loci detected in one strain only. Sequences alignment was performed by blastn.

**Supplementary Figure S5. CovR transcriptional regulation of cell-wall protein encoding genes.**

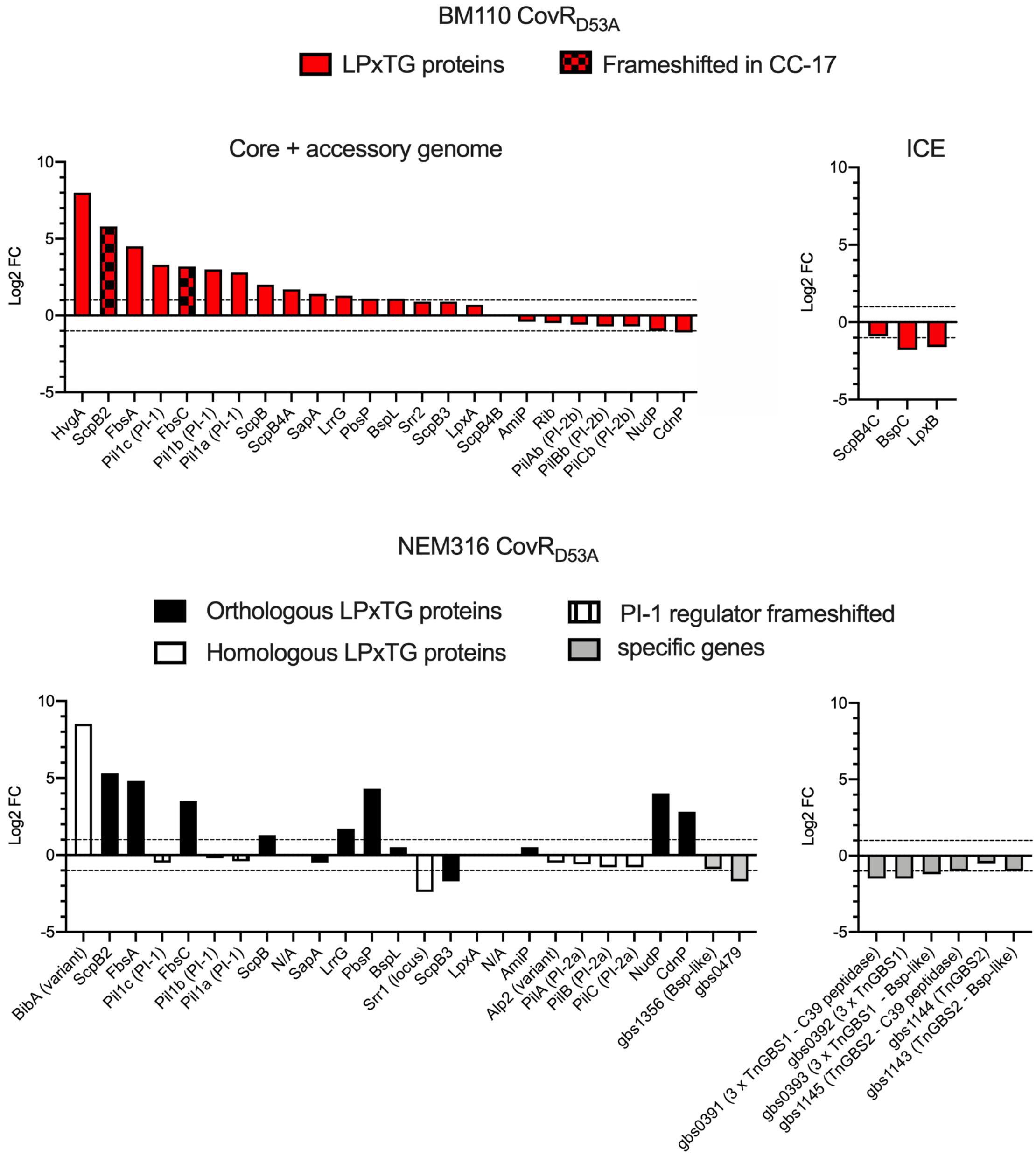


Histogram of RNA-seq fold changes (Log2 FC) in the CovR$_{D53A}$ mutants for the LPxTG encoding genes identified in the core and accessory genomes of BM110 (red upper panel) and NEM316 (black and white bottom panel). Columns in the NEM316 panel are color-coded following their similarities with BM110 proteins (black: orthologous genes; white: homologous genes including allelic variants and exclusive locus switching; grey: NEM316 specific genes; N/A: no homologue). The LPxTG encoded in integrative and conjugative elements (ICE) are grouped on the left due to the presence of several copies in NEM316 and the presence of several homologous but divergent genes between the two backgrounds. Translation of non-functional proteins due to mutations (frameshifts or internal stop) are indicated by hatched columns in BM110. Vertical lines on the three PI-1 proteins denoted the loss-of-function mutation (frameshift) of the PI-1 transcriptional activator in NEM316.

**Supplementary Figure S6. Comparison of CovR binding on the NEM316 and BM110 chromosomes.**

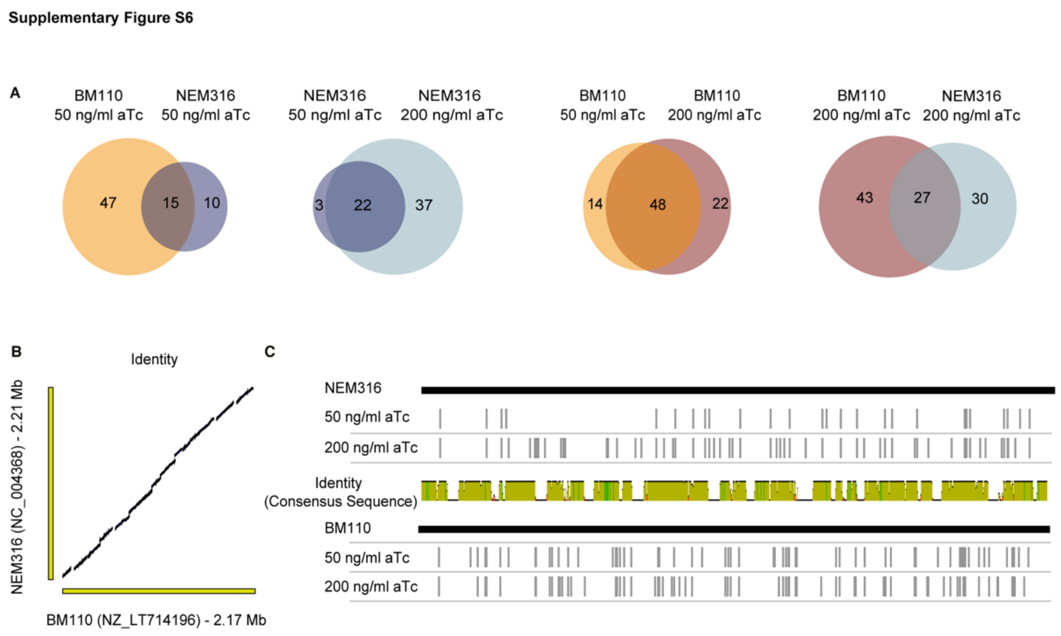


(A) Venn diagrams showing the overlaps between BM110 (orange and red) and NEM316 (blue and light blue) ChIP-seq experiments and between two levels of FLAG-CovR induction (50 and 200 ng/ml anhydro-tetracycline (aTc)). The numbers indicate significant peaks with a Fold Enrichment (FE) superior to 4 that are in common between two experiments or specific to one condition.

(B) Sequence comparison of the NEM316 and BM110 chromosomes. The two chromosomes are co-linear with vertical and horizontal gaps corresponding to strain-specific sequences, usually prophages and ICE.

(C) Linear representation of a consensus sequence between the NEM316 and BM110 chromosomes with the corresponding position of CovR binding sites identified by ChIP-seq. The consensus sequence represents nearly identical regions (yellow), regions with a high frequency of SNPs (green), and strain-specific sequences (white). The binding regions of CovR identified in each ChIP-seq experiment (NEM316 and BM110 at 50 and 200 ng/ml aTc induction) are symbolized by vertical grey lines.
