## Supplementary Tables for "The CovR regulatory network drives the evolution of Group B *Streptococcus* virulence": MVM et al sup table index.docx

**SUPPLEMENTARY TABLE TITLES** (excel files)

**Supplementary Table S1. RNA-seq analysis of *covR* mutants in BM110.**

Table S1A: Whole-genome analysis of RNA-seq in BM110 CovR_D53A_ and ∆*covR*

Table S1B: BM110 CovR_D53A_ up-regulated genes (Log2 FC > 1, adjusted p-value < 0.005)

Table S1C: BM110 CovR_D53A_ down-regulated genes (Log2 FC < -1, adjusted p-value < 0.005)

Table S1D: BM110 ∆ *covR* up-regulated genes (Log2 FC > 1, adjusted p-value < 0.005)

Table S1E: BM110 ∆ *covR* down-regulated genes (Log2 FC < -1, adjusted p-value < 0.005)

Table S1F: The BM110 *covR* regulon

**Supplementary Table S2. ChIP-seq analysis of CovR binding on the BM110 chromosome.**

Table S2A: CovR binding on the BM110 chromosome (with 50 ng/ml aTc, FE > 4 and IDR < 0.05)

Table S2B: Transcriptional changes associated to CovR binding

**Supplementary Table S3 : Transcriptional Start Sites (TSSs) in the BM110 chromosome.**

**Supplementary Table S4: Specificities of the CovR signaling pathway in the hypervirulent CC-17 lineage**

**Supplementary Table S5: RNA-seq analysis of the CovR_D53A_ mutant in NEM316.**

Table S5A: Whole-genome analysis of RNA-seq in NEM316 CovR_D53A_

Table S5B: Significantly and differentially transcribed genes (-1 > Log2 FC > 1, adjusted p-value < 0.005) in NEM316 CovR_D53A_

**Supplementary Table S6: RNA-seq analysis of the CovR core regulon in BM110 and NEM316.**

Table S6A. Whole RNA-seq data on the 1,716 orthologous genes in NEM316 and BM110

Table S6B. The CovR_D53A_ core regulon

Table S6C. The CovR_D53A_ strain-specific regulon

**Supplementary Table S7: LPxTG anchoring motif containing proteins in BM110**

**Supplementary Table S8: ChIP-seq analysis of CovR binding on the BM110 and NEM316 chromosomes at two levels of CovR induction**

Table S8A: CovR binding on the NEM316 chromosome with 50 ng/ml aTc and FE > 4

Table S8B: CovR binding on the NEM316 chromosome with 200 ng/ml aTc and FE > 4

Table S8C: CovR binding on the BM110 chromosome with 200 ng/ml aTc and FE > 4

**Supplementary Table S9: Comparative analysis of covR binding on the BM110 and NEM316 chromosomes**

Table S9A: Chromosomal CovR binding overlap between BM110 and NEM316

Table S9B: Sequences similarities between BM110 and NEM316 CovR binding regions (250 bp)

**Supplementary Table S10: IDR analysis of ChIP-seq experiments (FE > 1, IDR < 0.05)**

Table S10A: MACS2 and IDR analysis of ChIP-seq data (BM110 / 0 ng/ml aTc)

Table S10B: MACS2 and IDR analysis of ChIP-seq data (BM110 / 50 ng/ml aTc)

Table S10C: MACS2 and IDR analysis of ChIP-seq data (BM110 / 200 ng/ml aTc)

Table S10D: MACS2 and IDR analysis of ChIP-seq data (NEM316 / 0 ng/ml aTc)

Table S10E: MACS2 and IDR analysis of ChIP-seq data (NEM316 / 50 ng/ml aTc)

Table S10F: MACS2 and IDR analysis of ChIP-seq data (NEM316 / 200 ng/ml aTc)

**Supplementary Table S11: Strains, oligonucleotides, and plasmids**

Table S11A. Strains used in this study

Table S11B. Plasmids used in this study

Table S11C: Oligonucleotides sequences

Table S11D. Detailed plasmid construction
